# The nuclear receptor NR4A1 serves as a neutrophil-intrinsic regulator mitigating stroke severity

**DOI:** 10.1101/2025.09.19.677282

**Authors:** Jan-Kolja Strecker, Yannick Schmidt, Marie Liebmann, Carolin Walter, Stefan Roth, Hanna Aleth, Anna-Lena Börsch, Antje Schmidt-Pogoda, Carolin Beuker, Ayan Mohamud Yusuf, Julian Revenstorff, Collins Osei-Sarpong, Lisa Krafft, Anna-Sophie Schwarze, Michael Heming, Mathis Richter, Tanja Kuhlmann, Stephanie Hucke, Jan-Niklas Heming, Uwe Hansen, Nina Hagemann, Almke Bader, Jincheng Gao, Barbara Walzog, Daniela Maier-Begandt, Raphael Chevre, Eva-Maria Hanschmann, Martin Lücke, Clemens Sommer, Dirk M. Hermann, Johannes Roth, Christian Thomas, Gerd Meyer zu Hörste, Matthias Gunzer, Thomas Vogl, Alexander Zarbock, Frank Rosenbauer, Carlos Silvestre-Roig, Oliver Söhnlein, Luisa Klotz, Jens Minnerup

## Abstract

Ischemic stroke is accompanied by recruitment and activation of immune cells which play an important role in the progression of the brain damage. The nuclear receptor NR4A1 emerged as a key regulator within the inflammatory response of several immune diseases by regulating immune cell activation. In this study, we investigated the role of NR4A1 in the activation and recruitment of brain resident and peripheral immune cells after cerebral ischemia. Here, we show that NR4A1 mediates an anti-inflammatory and damage-limiting effect after stroke. This effect is largely mediated by neutrophil recruitment and importantly, NR4A1 activation with its ligand Cytosporone B improves functional outcome and reduces brain damage. Modulation of NR4A1 is therefore a promising therapeutic target for the treatment of the nuclear receptor NR4A1 in the activation and recruitment of peripheral and brain resident immune cells after cerebral ischemia and its consequences for stroke outcome. We demonstrate that NR4A1 ablation augments neutrophil activation and CNS recruitment within days after stroke thereby increasing infarct size, CNS inflammation, neuronal damage and deteriorating functional outcome. This effect is mediated via modulation of cell-intrinsic neutrophil function and maturation as illustrated by neutrophil-specific NR4A1 ablation and mixed bone-marrow chimera experiments. Notably, the NR4A1 agonist Cytosporone B reduced CNS neutrophil infiltration, infarct size and functional outcome after stroke in a bicentric preclinical stroke trial, demonstrating that NR4A1-mediated control of neutrophil reactivity is amenable to pharmacological modulation. In humans, NR4A1 expressing neutrophils are present in the peripheral blood of stroke patients and neutrophil NR4A1 expression correlates with improved long-term outcome after 3 months. Furthermore, NR4A1 expression in brain parenchyma neutrophils is negatively correlated with neuronal cell loss, illustrating a role of NR4A1 in regulating neutrophil mediated neuronal cell death in human stroke. Together our data reveal the nuclear factor NR4A1 as a brake of intrinsic neutrophil activity controlling neutrophil-mediated brain inflammation and neurotoxicity in stroke which may serve as a novel therapeutic target to limit inflammation-associated augmentation of ischemic damage after stroke.

## Introduction

Even after decades of research, stroke remains a leading cause of death and disability worldwide (Benjamin et al., 2017; Tsao et al., 2023). Intravenous thrombolysis and endovascular thrombectomy are landmark interventions, but their use is restricted by narrow therapeutic windows and contraindications, leaving most patients without access to reperfusion therapies (Powers et al., 2019). Cerebral infarction results from the sudden loss of blood flow to a brain region, causing severe oxygen and glucose deprivation. This metabolic stress triggers excitotoxic neurotransmitter release, mitochondrial dysfunction, and oxidative stress, leading to neuronal death via necrosis and apoptosis. Blood-brain barrier disruption further aggravates damage by allowing peripheral immune cell infiltration and amplifying local inflammation (Salaudeen et al., 2024). These processes create an irreversibly damaged infarct core and a surrounding penumbra of at-risk but potentially salvageable tissue (Ernime et al., 2021). In the acute phase, local inflammation accelerates brain injury through thromboinflammatory cascades that worsen barrier disruption and neuronal apoptosis (Denorme et al., 2022; Gullotta et al., 2023; Li et al., 2024). Neutrophils invade ischemic tissue within hours, releasing ROS, matrix metalloproteinases (MMPs), and neutrophil extracellular traps (NETs) that promote microvascular occlusion and neurotoxicity (Cai et al., 2020). NET density in thrombi correlates with thrombectomy resistance and poor outcomes, underscoring neutrophils as key regulators of stroke progression (Lapostolle et al., 2023). Recent studies further revealed neutrophil heterogeneity, identifying several functional subsets (Cai et al., 2020; Lapostolle et al., 2023). Nuclear receptors offer promising opportunities for modulating immune responses after stroke. These transcription factors regulate diverse processes including cell survival, metabolism, and stress adaptation, and many already have available ligands or antagonists (Jin et al., 2018; Shao et al., 2020). In particular, the NR4A family and NR4A1 (Nur77) in particular has emerged as a transcriptional checkpoint controlling myeloid cell activation. NR4A receptors are implicated in both immune- and non-immune– mediated diseases such as psoriasis, diabetes, rheumatoid arthritis, atherosclerosis, and cancer, acting via NF-κB regulation, metabolic reprogramming, and oxidative stress responses (Rahotra, 2015; Xiong et al., 2020; Crean & Murphy, 2021; Sheng et al., 2023). NR4A1 suppresses pro-inflammatory cytokines (IL-1β, IL-6, CXCL-8) in monocytes, macrophages, and T cells by inhibiting NF-κB (Hanna et al., 2011, 2012). Based on its role in inflammation and cerebral injury (Liebmann et al., 2018; Wang et al., 2018), we hypothesized that NR4A1 is a critical regulator in ischemic stroke. Using loss-of-function approaches in *Nr4a1*-deficient, chimeric, and cell-specific knockout mice, we identified infiltrating neutrophils as the main mediators of its protective effects. NR4A1 restricted neutrophil activation and recruitment to lesions, while pharmacological activation with the NR4A1 ligand Cytosporone B improved outcomes and reduced infarct size. Analyses of human brain tissue and blood confirmed NR4A1 expression in neutrophils after stroke, with expression levels correlating with patient outcomes.

## Results

### *Nr4a1* knock-out mice display greater infarct size accompanied by a significant increase in mature neutrophils within the ischemic brain

To determine the relevance of NR4A1 in ischemic stroke, we induced experimental stroke by middle cerebral artery occlusion of 30 min in *Nr4a1* knock-out versus wild type mice (Fig 1a). As depicted in figure 1b, 72h post ischemia, infarct volume was significantly larger in *Nr4a1* knock-out animals, and this was accompanied by increased locomotor deficits (Fig 1c). Histological evaluation of infarcted brain areas using TUNEL-staining (TdT-mediated dUTP-biotin nick end labeling) revealed increased neuronal cell death in *Nr4a1* knock-out mice (Fig 1d). After experimental stroke, NR4A1 expression on mRNA and protein level substantially increased within the infarcted area with a peak at 72h h (Suppl Fig 1a), suggesting that it is expressed by invading immune cells. Flow cytometric quantification of CNS immune cell infiltration and activation after stroke induction did not reveal relevant changes in microglial cells, macrophages, and T cells (Fig 1e, Suppl Fig 1b), but a strong increase in neutrophils in the CNS of *Nr4a1* knock-out mice as compared to their wildtype counterparts (Fig 1f, g). Notably, infarct size correlated negatively with the proportion of NR4A1 expressing mature but not immature neutrophils in the peripheral blood (Fig 1h, Suppl Fig 1c). Further characterization of CNS neutrophils pointed to a predominant presence of mature (i.e. segmented) neutrophils in both wild type and knock-out animals, while the difference between wild type and knock-out animals was driven by a predominant increase in ROS-producing (HNE^+^) mature neutrophils, pointing towards an enhanced activity (Fig 1i). Together, these data point towards an unexpected role of NR4A1 in neutrophils with implications for stroke severity. Worsened functional outcome in knock-out animals at 72 hours but not at 24 or 48 hours further suggests a delayed protective effect of NR4A1 after the very early acute phase of cerebral ischemia. Of note, we could exclude that the observed larger infarcts in *Nr4a1* knock-out animals and the increase in neutrophil numbers were mediated by changes in cerebral vessel occlusion, as there was no difference in fibrinogen deposition within cerebral vessels (Fig 1j). Neutrophil phagocytosis by microglial cells was not altered in *Nr4a1* knock-out as compared to wild type mice (Suppl. Fig 1d). Moreover, neutrophil apoptosis rates within the CNS during stroke were not significantly altered in *Nr4a1* knock-out animals, indicating that increased neutrophil numbers in the CNS of *Nr4a1* knock-out mice during stroke are not due to an increased neutrophil live span (Suppl. Fig 1e).

**Figure 1.**
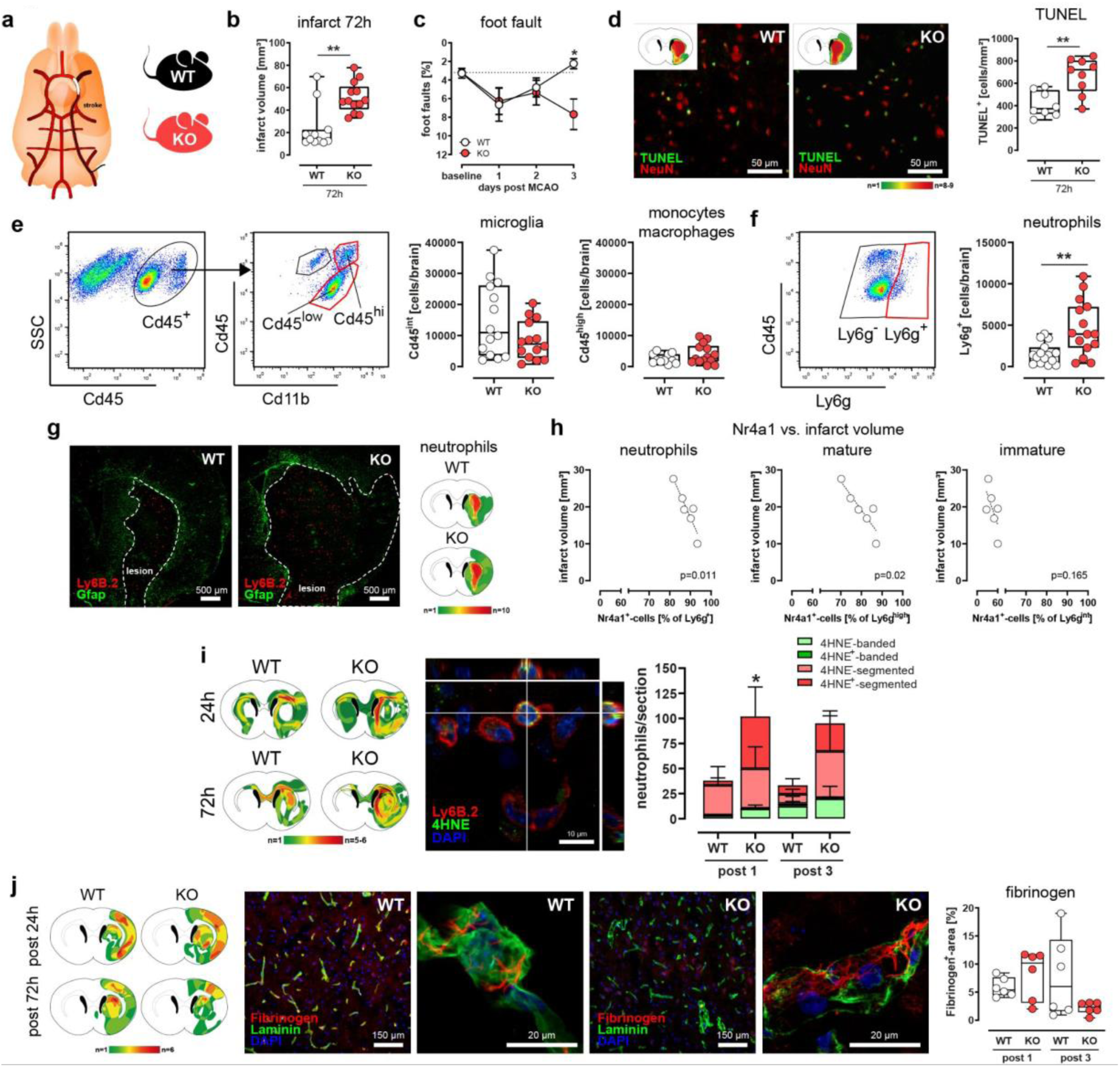
Nr4a1 exerts protective effects in cerebral ischemia. **a,** Experimental cerebral ischemia was induced in wild-type and *Nr4a1*-deficient mice. **b,** Mean infarct volumes were calculated from cryosections of wild type and *Nr4a1*-deficient mice at 72h after reperfusion (WT n=11, KO n=13; \*\**P*=0.002, two-tailed *t*-test). Median cell counts are shown as box plots. Lines within the boxes denote medians. **c,** Locomotor function was assessed with the foot-fault test (WT n=14, KO n=17; \**P*=0.021 post 3, two-way ANOVA/mixed model) prior to MCAO and on day 1, 2 and 3 post reperfusion. **d,** Detection of damaged/apoptotic neurons was performed using TUNEL/NeuN double-staining on cryosections at bregma 0-1 mm. Quantification was performed within the ischemic striatum and cortex. Median cell counts are presented as box plots. The lines inside the boxes denote medians (WT n=8; KO n=9, \*\**P*=0.002, two-tailed *t-*test). **e,** Gating strategy for sorting and quantification of intracerebral Cd45^low^/Cd11b^+^-microglia and Cd45^high^/Cd11b^+^-myeloid cells. **f,** Neutrophil quantification was performed by FACS-analysis counting Cd45^+^/Ly6g^int^ and Cd45^+^/Ly6g^high^-cells. Median cell counts are presented as box plots. The lines inside the boxes denote the medians (WT n=15; KO n=15, \*\**P*=0.002, *t*-test). **g,** Immunohistological analysis of neutrophil infiltration (Ly6b.2, red) in relation to the astroglia scar (GFAP, green) in wild type and *Nr4a1*-deficient mice. Neutrophil heatmaps were generated from brains post 72h (n=10 each). **h,** Correlation between individual infarct volumes of wild type mice subjected to MCAO (post 72h) and the number of peripheral NR4A1-expressing Ly6g^+^-neutrophils (post 24h, \**P*=0.011, linear regression) as well as number and NR4A1-expression intensity of the circulating subpopulation of mature Ly6g^high^-neutrophils (post 24h, \**P*=0.02, linear regression). **i,** Quantification of 4HNE^−^- and 4HNE^+^-immature (banded nucleus) and mature (segmented nucleus) neutrophils (1d post MCAO each group n=5; 3d post MCAO WT n=5; KO n=6; *P=0.0372; two-way ANOVA). **j,** Quantification and heat map generation of fibrin deposition was performed on immunohistochemically stained coronal sections (fibrinogen, red; vascular marker laminin green) from brains at post 24h and 72 h (n=6 per group and time-point). Quantification and heatmap generation of lipid oxidation products (4-HNE) were performed on immunohistochemically stained brain sections.

### *Nr4a1* expression in neutrophils determines stroke severity and infarct size

We next performed a series of experiments to determine which cell types are responsible for NR4A1-mediated modulation of infarct size and stroke severity, as NR4A1 is expressed in a number of immune cell types and also CNS resident cells (Suppl. Fig 1 f-h). First, bone-marrow chimeras revealed that cells of hematopoietic origin are responsible for the increase in stroke severity by lack of *Nr4a1*, as transfer of *Nr4a1*-deficient bone marrow into wild type recipients resulted in an increase in infarct volume, decreased motor function, and neuronal cell death (Fig 2a-c, Suppl Fig 2a+b). Adoptive transfer of either wildtype T cells or *Nr4a1*-deficient T cells to *Rag-1*-deficient mice (devoid of T and B cells, Fig 2d, Suppl Fig 2c) showed no differences with respect to locomotor function outcome (Fig 2e) and infarct volume (Fig 2f), indicating that the effect of NR4A1 in stroke is not mediated via T cells, which had been shown by us before in the context of experimental autoimmune encephalomyelitis (Liebmann et al., 2018). In this line, mice lacking *Nr4a1* cell-specific in T cells (Fig 2g) showed no altered functional outcome (Fig 2h) and infarct volume (Fig 2i) when compared to the Cre negative control group. Given the increased number of neutrophils in the infarcted brains of *Nr4a1* deficient mice, we generated conditional mice lacking *Nr4a1* specifically in neutrophils upon Cre-mediated *Nr4a1* ablation in *Ly6g*-Cre expressing cells (Fig 2k). Notably, neutrophil-specific *Nr4a1* ablation completely recapitulated the phenotype of *Nr4a1* full knock-out mice, illustrated by decreased motor function (Fig 2k) and increased infarct volume (Fig 2l), pointing towards a key role of NR4A1 in neutrophils for increased stroke severity.

**Figure 2.**
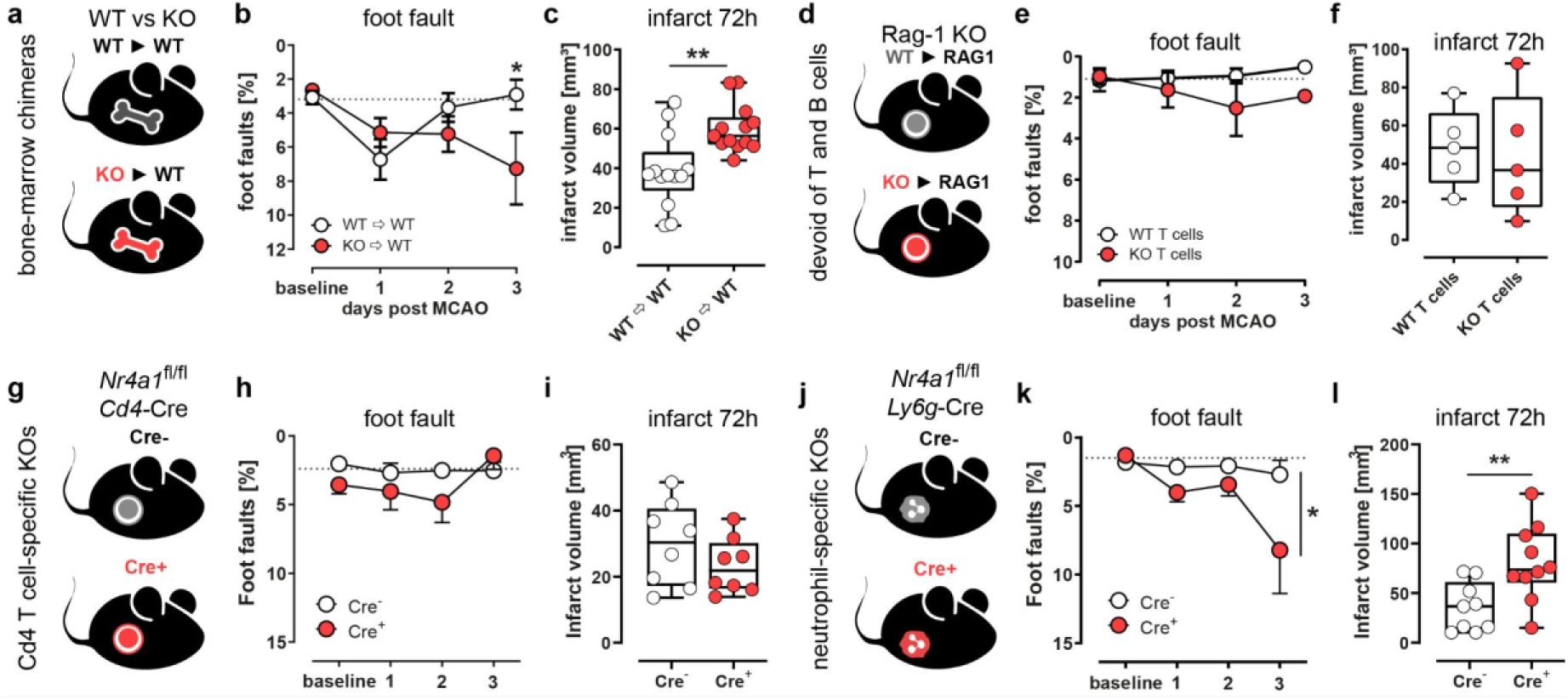
Nr4a1 protective effects are mediated by neutrophil granulocytes. **a,** Schematic depiction of the chimerism approach. Cd45.1 congenic wild-type mice (WT) or Cd45.2 *Nr4a1*-deficient (KO) were sublethally irradiated and reconstituted with either congenic wild-type (WT Cd45.2►WT Cd45.1) or *Nr4a1*-deficient (KO Cd45.2►WT Cd45.1) bone-marrow cells (WT Cd45.2►KO; Cd45.1). **b,** Foot-Fault tests were performed prior to MCAO (baseline) and 24h, 48h, and 72h after MCAO (each group n=13; \**P*<0.05, ANOVA). **c,** Mean infarct volumes from cryosections of WT►WT and KO►WT mice at 72h after reperfusion. Lines within the boxes denote medians (each group n=13; \**P*=0.0018, two-tailed *t*-test). **d,** Schematic depiction of adoptive T cell transfer. Either CD3^+^/WT T cells or CD3^+^/*Nr4a1*-deficient T cells were intravenously administered to *Rag-1*-deficient mice. **e,** *Rag-1*-deficient mice were subjected to foot-fault testing prior to and 24h, 48h and 72h after MCAO. **f,** Infarct volumes were compared between *Rag-1*-deficient mice which received either WT- or *Nr4a1*-deficient T cells. **g,** Conditional knock-out mice lacking *Nr4a1* in T cells were generated by crossing *Nr4a1*fl/fl-mice with mice expressing a Cre-recombinase under the control of *Cd4*-promotor. **h,** Foot-Fault tests were performed prior to MCAO (baseline) and 24h, 48h, and 72h after MCAO (Cre^+^ n=8; Cre^−^-n=8) **i,** Mean infarct volumes were calculated from cryosections of Cre^+^- and Cre^−^-mice at 72h after reperfusion. Median cell counts are shown as box plots. Lines within the boxes denote medians (Cre^+^ n=8; Cre^−^ n=8)**. j,** Conditional knock-out mice lacking *Nr4a1* in mature neutrophils were generated by crossing *Nr4a1*^fl/fl^-mice with mice expressing a Cre-recombinase/tdTomato reporter under the control of *Ly6g*-promotor. **k,** Foot-Fault tests were performed prior to MCAO (baseline) and 24h, 48h, and 72h after MCAO (Cre^+^ n=12; Cre^−^ n=10, \**P*=0.0132, two-way ANOVA/mixed model) **l,** Mean infarct volumes were calculated from cryosections of Cre^−^- and Cre^+^-mice at 72h after reperfusion. Median cell counts are shown as box plots. Lines within the boxes denote medians (Cre^+^ n=12; Cre^−^ n=10; \*\**P*=0.0083, *t*-test).

### NR4A1 mediated neutrophil-intrinsic effects modulate stroke severity

Given the central role of neutrophils for mediating NR4A1 effects in the context of stroke, we next addressed whether NR4A1 rather affects neutrophil recruitment or directly modulates intrinsic neutrophil activity. First, we investigated whether healthy animals already show differences in peripheral blood neutrophil count or activation. Flow cytometry data showed no significant difference with respect to the number of circulating immature and mature neutrophils between WT and KO mice (Fig. 3a). Analysis of the activation and maturation markers CD182, Ly6B.2, CD14 (Sas et al., 2020) and CD133 (Kim et al., 2017) also revealed no differences between the examined genotypes under steady state condition (Fig. 3b). Electron microscopy was further employed to assess potential morphological alterations in bone marrow-derived neutrophils from healthy wild type and *Nr4a1*-deficient mice but no obvious differences were observed between the groups (Fig. 3c). Next, we determined the expression levels of neutrophil-relevant chemokines within the CNS of *Nr4a1* knock-out and wild type mice during stroke; however, we did not detect significant differences, in particular for *Ccl2*, *Cxcl1* and *Cxcl2* (Fig. 3d, suppl. Fig. 2e), indicating that recruitment to the brain of *Nr4a1* deficient mice was not driven by enhanced local chemokine signaling. Furthermore, during stroke, we observed reduced levels of mature neutrophils in the peripheral compartments, especially blood, bone-marrow and spleen, whereas frequencies of mature neutrophils were significantly higher in the CNS of *Nr4a1* knockout animals, pointing towards an increased neutrophil egress and maturation (Fig. 3e). Next, we generated mixed bone-marrow chimeras by transplantation of an equal ratio of GFP-labeled CD45.2 wild type and unlabeled CD45.2 *Nr4a1* knock-out bone-marrow into wild type CD45.1 mice and subjected these animals to ischemic stroke (Fig. 3f, Suppl. Fig. 2a). At baseline, wild type and knock-out neutrophil ratios were equal in the peripheral blood, but at days 1 and 2 post stroke induction the proportion of knock-out neutrophils was significantly increased (Fig. 3f), and after 72h the proportion of knock-out neutrophils was likewise elevated in the CNS of stroke mice (Fig 3h). This indicates that *Nr4a1* ablation in neutrophils facilitates their egress from the bone marrow and subsequent migration into the CNS in the context of stroke. To further elucidate the cell-intrinsic effects of NR4A1 on neutrophils, we performed a set of *in-vitro* experiments using HoxB8-derived *Nr4a1* knock-out and wild type neutrophils as an established model system to study neutrophil functions *in vitro*. First of all, apoptosis rates were not altered, indicating that NR4A1 does not alter neutrophil live span per se (Suppl. Fig 3a). Further, ablation of *Nr4a1* did not alter neutrophil adhesion and migration activities (Suppl. Fig 3b-d). However, we observed significant alterations of neutrophil metabolic activity, as illustrated by an increased respiratory capacity of *Nr4a1* knock-out neutrophils under inflammatory conditions (Fig 3i). Furthermore, *Nr4a1* knock-out neutrophils exhibited increased ROS activity (Fig 3j). Bulk RNA-sequencing of *Nr4a1* knock-out versus wild type HoxB8 neutrophils at earlier versus later stages of differentiation revealed an upregulation of biological processes related to stress response, cellular activation and cell division (Fig 3k). Heatmap visualization of gene set enrichment analysis revealed an enrichment of *Nr4a1*-deficient gene signatures associated with oxidative response, neutrophil maturation and activation, NETosis, phagocytosis and chemotaxis, pointing towards an enhanced activation state of neutrophils in the absence of NR4A1 (Fig 3l).

**Figure 3.**
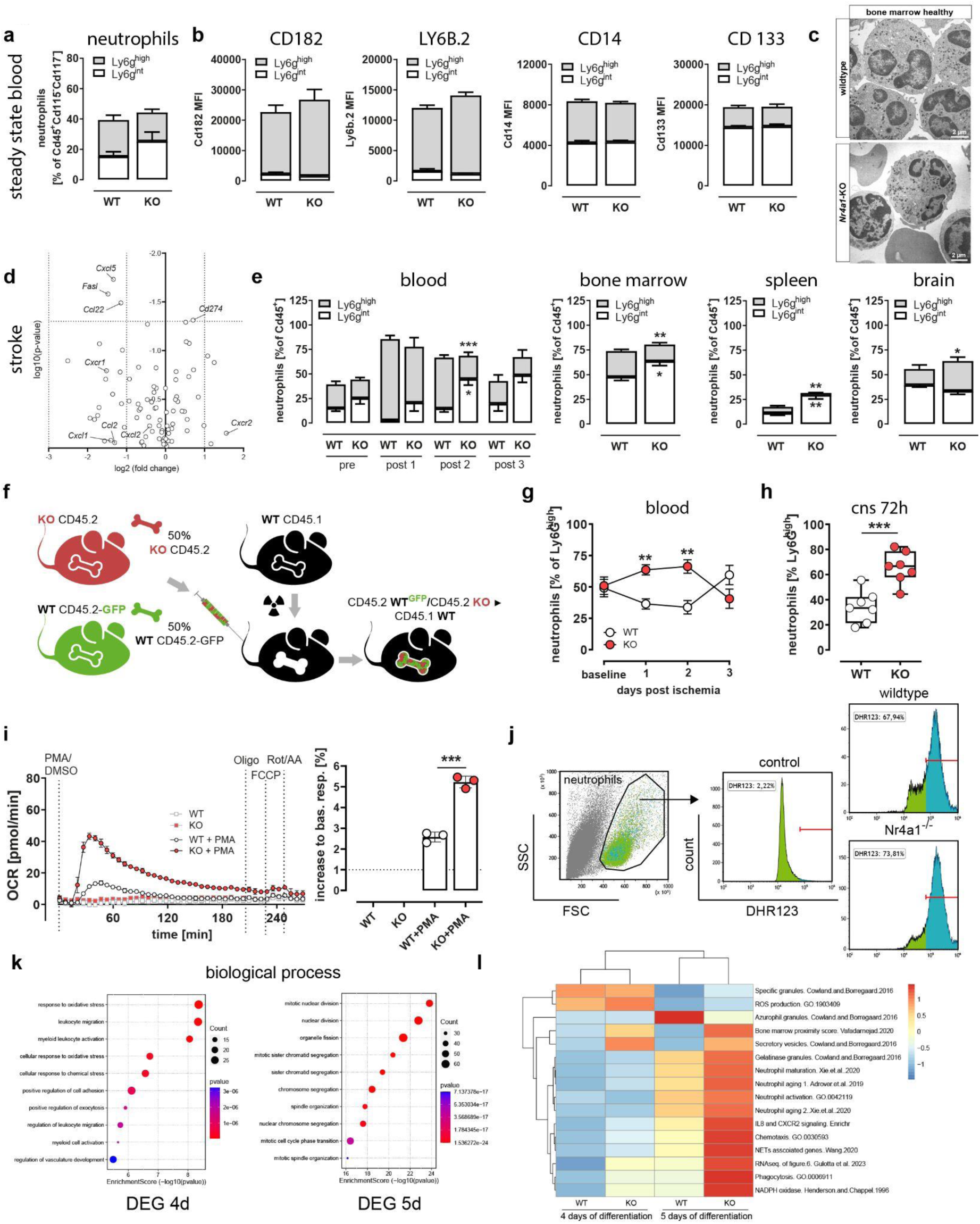
Nr4a1 regulates neutrophil activation after experimental stroke. **a,** FACS analysis of circulating neutrophils was performed with peripheral blood of healthy wild type and *Nr4a1* deficient mice (WT n=6; KO n=6). **b,** FACS analysis of neutrophil expression of maturation and activation marker CD182, LY6.B2, CD14, CD133 was performed using peripheral blood of healthy wild type and *Nr4a1* deficient mice (WT n=6; KO n=6). **c,** Ultrastructure analysis was performed on wild type and *Nr4a1* deficient bone-marrow cells using electron microscopy. **d,** RT-PCR-expression analysis of inflammation-related genes in WT and KO mice 24h after MCAO. mRNA was harvested from whole-brain lysates (WT n=4; KO n=5). **e,** Quantification of immature and mature peripheral neutrophils was performed over the course of 3 days post MCAO by FACS (**\***\**P*=0.005; two-way ANOVA). Quantification of bone marrow neutrophils 72h after MCAO (immature \**P*=0.0243, mature \*\**P*=0.0072; unpaired two-tailed *t*-test). Quantification of splenic neutrophils 72h after MCAO. (immature ** *P*=0.0043, mature ** *P*=0.0042; unpaired two-tailed *t*-test). Quantification of intracerebral neutrophils 72h after MCAO. (mature ** *P*=0.039; unpaired two-tailed *t*-test). Median cell counts are shown as box plots. Lines within the boxes denote medians (WT n=5; KO n=5). **f,** Mixed-bone marrow chimeras were generated by injecting 50% WT and 50% KO-bone-marrow into irradiated congenic Cd45.1 mice. **g,** FACS-quantification of WT and KO peripheral neutrophils was performed prior to MCAO and over the course of 3 days post MCAO. Data are presented as mean ± s.e.m, (n=7; post 1d **\***\**P*=0.0021; post 2d **\***\**P*=0.0045; two-way ANOVA). **h,** FACS-analysis of intracerebral WT and KO neutrophils 72h after MCAO. Median cell counts are shown as box plots. Lines within the boxes denote medians (n=7; \*\**P*=0.0006; unpaired two-tailed *t*-test). **i,** Oxygen consumption rate (OCR) of wild type and *Nr4a1*-deficient HoxB8-immortalized neutrophil granulocytes were determined following 5 days of differentiation. Cells were stimulated either with or without 600 nM phorbol-12-myristat-13-acetat (PMA, Sigma-Aldrich) for 3h. **j,** ROS-generation of HoxB8-neutrophils was evaluated by FACS DHR123-assay. **k,** Dotplot of the ten most enriched gene-sets for GO Biological Processes. X-axis shows −log10 p value; the dot size represents the number of genes in the significant DE gene list associated with the GO term. **l,** Heatmap showing the expression pattern of gene sets in wild-type and *Nr4a1*-deficient HoxB8-neutrophils after 4d and 5d of differentiation.

### NR4A1 shows no effect on neutrophils under physiological conditions but attenuates neutrophil activation after experimental stroke

We next sought to further elucidate the effects of *Nr4a1* deficient neutrophils within the CNS after stroke. We performed single-cell RNA sequencing of neutrophils isolated from the CNS of stroke animals, alongside neutrophils from blood and bone marrow for comparison (Fig 4a). Cluster analysis identified distinct populations in blood, bone marrow, and CNS. Due to the lower cell number, CNS neutrophils were subdivided into three clusters (Suppl. figure 4a+b). Comparison of wild type and *Nr4a1* knock-out neutrophils revealed no relevant differences in blood clusters (Fig. 4b) and only a slight increase of knock-out cells in bone marrow cluster 5 (Fig. 4c). In contrast, CNS populations showed pronounced changes, with strong enrichment of *Nr4a1* knock-out neutrophils in cluster 1, accompanied by a decrease in clusters 0 and 2 (Fig. 4d). Enrichment analysis of custom neutrophil signatures showed that CNS cluster 0 is enriched in more mature neutrophils characterized by extracellular matrix damage, sterile hyperinflammation, and necroptosis, whereas cluster 1 represents a classic acute effector profile associated with early injury, including NETosis, phagocytosis, and degranulation (Suppl. figure 4c+d). Direct transcriptomic comparison of wild type and *Nr4a1* knock-out neutrophils (Fig. 4e) revealed upregulation of genes related to chemotaxis and recruitment (*Cxcl2*, *Cxcl3*, *Cxcrs*), degranulation and tissue injury (*Elane*, *Prtn3*, *Mpo*, *Mmp9*, *Plek*), NETosis, inflammatory signaling (*Il1a*, *Il1b*, *Tnf*, *Nfkbia*, *Nfkbiz*), and metabolic reprogramming and stress response (*Hilpda*, *Fabp5*, *Apoe*, *Acod1*, *Sfxn5*, *Slc7a11*, *Hspa1a*). Consistently, GO term analysis confirmed this *Nr4a1* knock-out phenotype, showing upregulation of pathways related to cytokine production, intracellular stress responses, migration and chemotaxis, and neuronal damage (Fig. 4f). Additional enrichment analysis using a custom neutrophil signature database supported these findings, demonstrating significant enrichment for maturation, PAMP recognition, chemotaxis, inflammatory signaling, and cell death (Fig. 4g). Together, these results demonstrate that *Nr4a1* deletion in neutrophils promotes their migration into the CNS, enhances their pro-inflammatory state, and increases their tissue-destructive potential. Notably, histologic analysis revealed that a substantial proportion of neutrophils was found in close proximity to neurons, with the prevalence of such interactions increasing along with neutrophil maturation (Fig. 4h, i). Hypersegmented neutrophils exhibited a significant increase in neutrophil - neuron interactions in both global *Nr4a1*-knockout mice (Fig. 4j) and neutrophil-specific knockout animals (Fig. 4k), indicating that NR4A1 could modulate and limit neutrophil attraction to distressed neurons. As this observation was novel and, to our knowledge, not previously reported, we next examined additional experimental stroke models to assess whether direct neutrophil - neuron interactions could be a common cellular response following stroke. Indeed, these interactions were also observed in *Ly6g*-reporter mice, in a rat model of tMCAO, and even in long-term brains subjected to photothrombotic stroke up to 49 days post-ischemia (Suppl. Fig. 4e-h). These findings suggest that neutrophil - neuron interaction may represent a common pathomechanism in cerebral ischemia.

**Figure 4:**
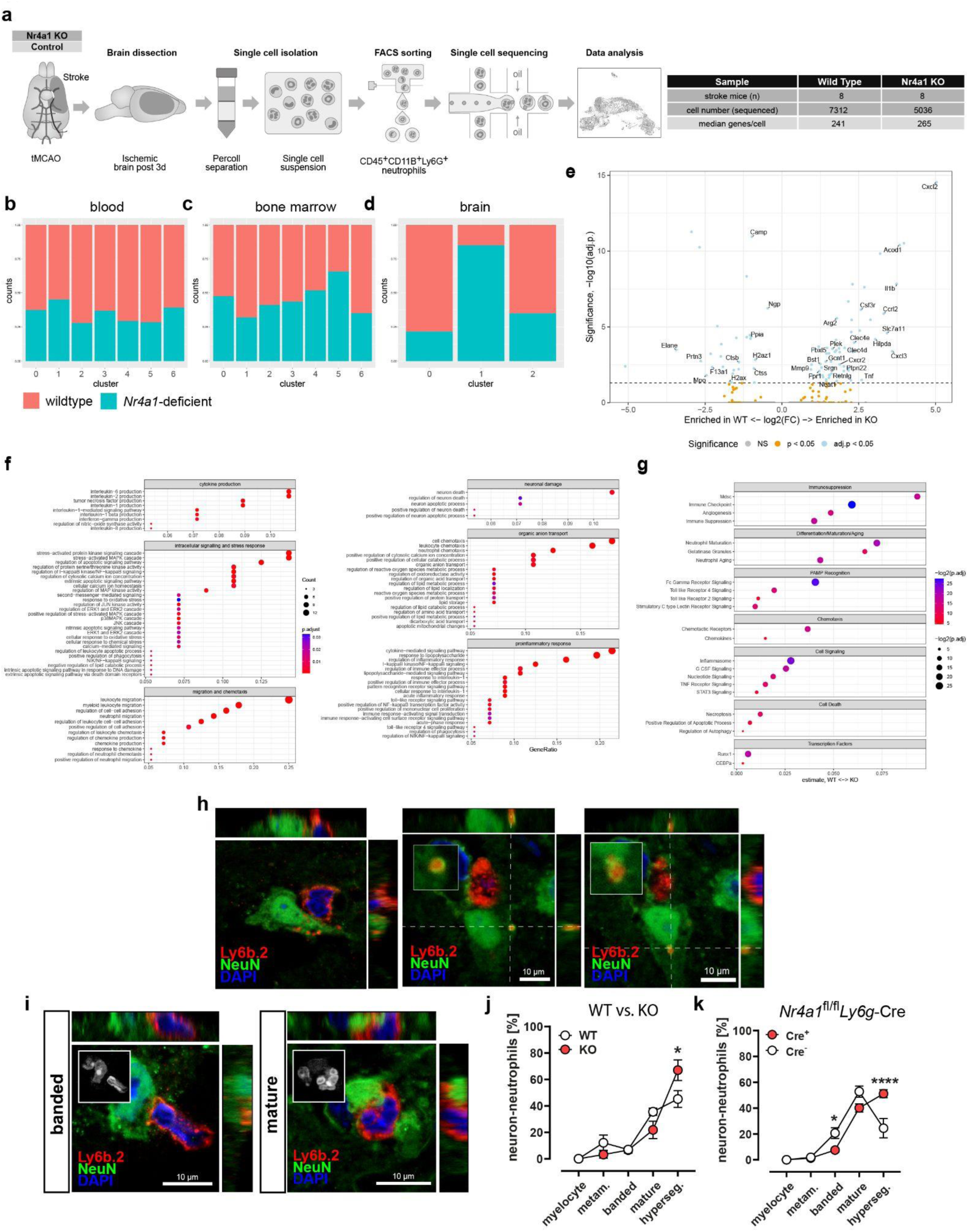
Single-cell transcriptomics analysis. **a,** Wild type (n=8) and *Nr4a1*-deficient mice (n=8) were subjected to transient middle cerebral artery occlusion (tMCAO). After 3 days of reperfusion brains were dissected, mechanically dissociated, and subjected to Percoll gradient centrifugation for single-cell isolation at 4 °C. CD45⁺CD11B⁺LY6G^+^-neutrophils were sorted by fluorescence-activated cell sorting (FACS) and processed for single-cell RNA sequencing. Data analysis was performed on tagged and pooled samples. Summary of sample characteristics is shown in the table, including total number of sequenced cells and median number of detected genes per cell. **b,** Relative proportions of wild type (red) and *Nr4a1*-deficient (blue) cell counts per cluster for blood, **c,** bone marrow and **d,** CNS. Due to differences in preprocessing, the CNS cell population was analyzed separately. **e,** Volcano plot of DEGs (differentially expressed genes) between wild type and *Nr4a1*-deficient CNS cell populations, displaying significantly down- or upregulated neutrophil related genes (adj. p<0.05, blue). **f,** Enrichment analysis of GO (gene ontology) BP (biological processes) terms for DEGs upregulated in CNS *Nr4a1*-deficient cells, compared to CNS wild type (adjusted p-value<0.05), with gene count and gene ratio per term (defined as relevant genes per GO BP term, divided by number of input genes). All terms were filtered and grouped into categories of interest. **g,** Enrichment analysis of a custom neutrophil signature database between CNS *Nr4a1*-deficient and CNS wild type cells, grouped by biological categories. Enrichment scores per cell were determined with the ESCAPE/UCell pipeline, and differences between conditions KO and wld type cell sets calculated with ANOVA and Tukey tests (adjusted p-value<0.05). **h,** Immunofluorescence photographs depicting neutrophils in close contact to neurons and neutrophil-specific Ly6b.2-signal within the damaged neurons. **i,** The degree of neutrophil maturation was determined at high magnification based on DAPI-stained nuclear morphology (box insert) and defined as banded, segmented or hypersegmented. **j,** Quantification and characterization of neutrophils attached to neurons depending on maturation grade in wild type versus *Nr4a1*-deficient mice (wild type n=6; *Nr4a1*-deficient n=9; *P=0.0118, two-way ANOVA). **k,** Neutrophil-neuron interactions were quantified in mature neutrophil cell-specific *Ly6g*-Cre mice (*Ly6g*-Cre+ n=6; Ly6g-Cre-n=5; **P=0.0361, two-way ANOVA).

### Pharmacological activation of NR4A1 via Cytosporone B in neutrophils ameliorates stroke severity

Based on our results so far, pointing towards a key role of NR4A1 for controlling neutrophil activation thus limiting their detrimental effects on infarct volume and CNS inflammation, we tested whether pharmacologic activation of NR4A1 using the agonist Cytosporone B can ameliorate stroke severity by modulation of neutrophil activity. To this end, mice were treated with Cytosporone B once (13 mg/kg) 3h after stroke induction. As depicted in Figure 5a, NR4A1 activation reduced the infarct volume and consecutively the stroke-associated decrease in motor function (Fig. 5b). Accordingly, we observed a significant reduction of CNS neutrophils in Cytosporone B treated animals (Fig 5c). Consequently, the percentage of interactions between mature neutrophils and CNS neurons was significantly reduced (Fig. 5d), and overall, we observed a significant preservation of neurons in the cortex of Cytosporone B treated animals (Fig. 5e). In *Nr4a1* knock-out animals, Cytosporone B treatment did not affect infarct volume and decrease in motor function, thus demonstrating that these effects are indeed mediated via NR4A1 (Fig. 5f).

**Figure 5.**
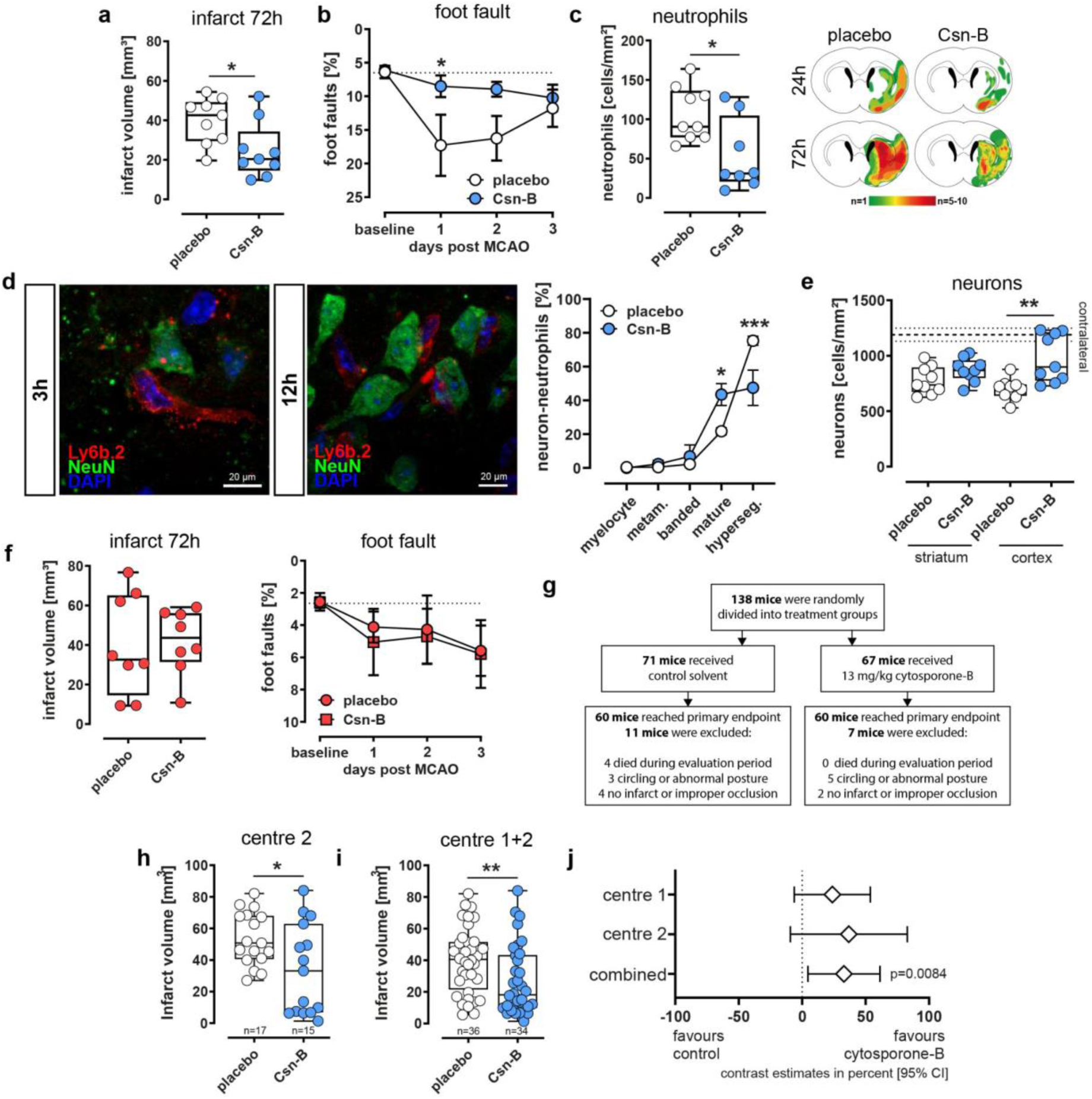
NR4A1 activation exerts protective effects in cerebral ischemia. **a,** Mean infarct volumes were calculated from toluidine stained coronal sections (each group n=9; \**P*=0.0121, *t*-test). **b,** Foot-Fault tests (Csn-B 3h post MCAO) were performed prior to (baseline) and 24h, 48h, and 72h after MCAO (placebo n=8; Csn-B n=10; \**P*<0.028, two-way ANOVA). **c,** Detection of intracerebral neutrophils. Median cell counts are presented as box plots. The lines inside the boxes denote medians (placebo n=8; Csn-B n=8, *P*=0.0161; two-tailed Mann-Whitney test). Generation of heatmaps (24h and 72h after MCAO) and quantification (72h after MCAO) of neutrophil presence was performed on immunohistochemically stained coronal slices at bregma 0-1 mm. **d,** Neutrophil-neuron interactions were quantified in placebo and Csn-B-treated mice (placebo n=4; Csn-B n=4, male; post 2d \**P*=0.0118, post 3d \*\**P*=0.0010, two-way ANOVA). The left panel shows neutrophil-neuron contact in Csn-B treated mice (Csn-B applied 3h and 12h post-stroke onset). **e,** Neurons were quantified on fluorescence stained sections within the ipsilateral striatum and cortex (each group n=9; \*\**P*<0.0027, unpaired two-sided *t-*test) of placebo and Csn-B treated mice (3h post MCAO). **f,** *Nr4a1*-deficient mice were treated with either placebo or Csn-B (13 mg/kg) 3h after MCAO. Foot-Fault tests were performed prior to MCAO and 24h, 48h, and 72h after MCAO (placebo n=5; Csn-B n=5). **g,** Design of the bi-centric study. Mice were treated either with 13mg/kg μg of Cytosporone B or placebo 3h after stroke. Randomization, distribution of Cytosporone B and blinded analysis of infarct volumes were performed in Muenster. **h,** Mean infarct volumes of placebo or Csn-B-treated mice generated at centre 2 were calculated at centre 1 from toluidine stained coronal sections (placebo female n=10, male n=7; Csn-B female n=8, male n=7; \**P*=0.0223, unpaired two-tailed *t*-test). **i,** Combined mean infarct volumes generated at centre 1 and centre 2 from coronal sections (placebo n=36; Csn-B n=34; \*\**P*=0.0084, Mann-Whitney test). **j,** Bicentric study contrast estimates for centre 1, centre 2 and combined for both centres.

To meet the increasing demand for preclinical studies in stroke research, we decided to conduct a randomized bicentric and pre-registered study on the efficacy of Cytosporone B and performed a bicentric blinded trial (Fig. 5g). As our initial experiments were performed in male mice only, we included an additional study in female mice at our site (centre 1, UKM). At centre 2 (UKE), 40 mice (20 males and 20 females; 10 placebo and 10 Csn-B each) were subjected to MCAO and treated with either placebo or Csn-B 3h after reperfusion. All mice were randomly assigned in a 1:1 ratio to receive either solvent or Csn-B. As depicted in Figure 5h-i, this bicentric study could corroborate our previous results, thus strengthening our concept of NR4A1 in neutrophils as a pharmacological target to limit inflammation-induced increases in stroke volume and consecutive stroke severity (Fig 5j). Together, our data provide evidence that NR4A1 in neutrophils serves as a cell-intrinsic brake of neutrophil activation and is a promising target for modulating neutrophil mediated augmentation of stroke severity in the early phase after stroke.

### Human blood and CNS neutrophils express NR4A1 and its expression correlates with long-term outcome after stroke

Next, we sought to validate our findings from the rodent experiments within human pathophysiology. We first characterized intracerebral neutrophils with respect to NR4A1 expression, maturation status, and neuronal interactions using both fresh-frozen and FFPE sections. Histological analysis revealed that, following ischemic stroke, immigrated neutrophils expressed NR4A1 (Fig. 6a) and were observed in close contact with neuronal proteins (Fig. 6b), and numerous immigrated cells were found adjacent or in contact with MAP2-positive dendrites and neurons (Fig. 6c). We then examined whether their maturation state influenced the interaction with neurons. On average, neutrophils found within the ischemic area during the subacute phase (day 5) showed a higher maturation grade than those present in the later starting chronic phase (day 15; Fig. 6d). In the next set of experiments, we strove to elucidate the potential translational relevance of our findings. First, in a monocentric human stroke cohort comprising 71 stroke patients we determined the expression of NR4A1 in peripheral blood neutrophils by flow cytometry in the acute phase (i.e. 48h after symptom onset) and correlated its expression with the neurological outcome at three months (modified Rankin scale, mRS) (Fig 6e+f). As expected, we observed that the initial clinical manifestation (measured by the NIHSS score) positively correlated with increased long-term disability (mRS). Notably, we could show that the percentage of NR4A1 expressing neutrophils negatively correlated with long-term disability (Fig 6g-i), therefore underlining the concept that pharmacological modulation of NR4A1 in neutrophils represents a promising therapeutic target to limit inflammation-induced augmentation of stroke severity.

**Figure 6.**
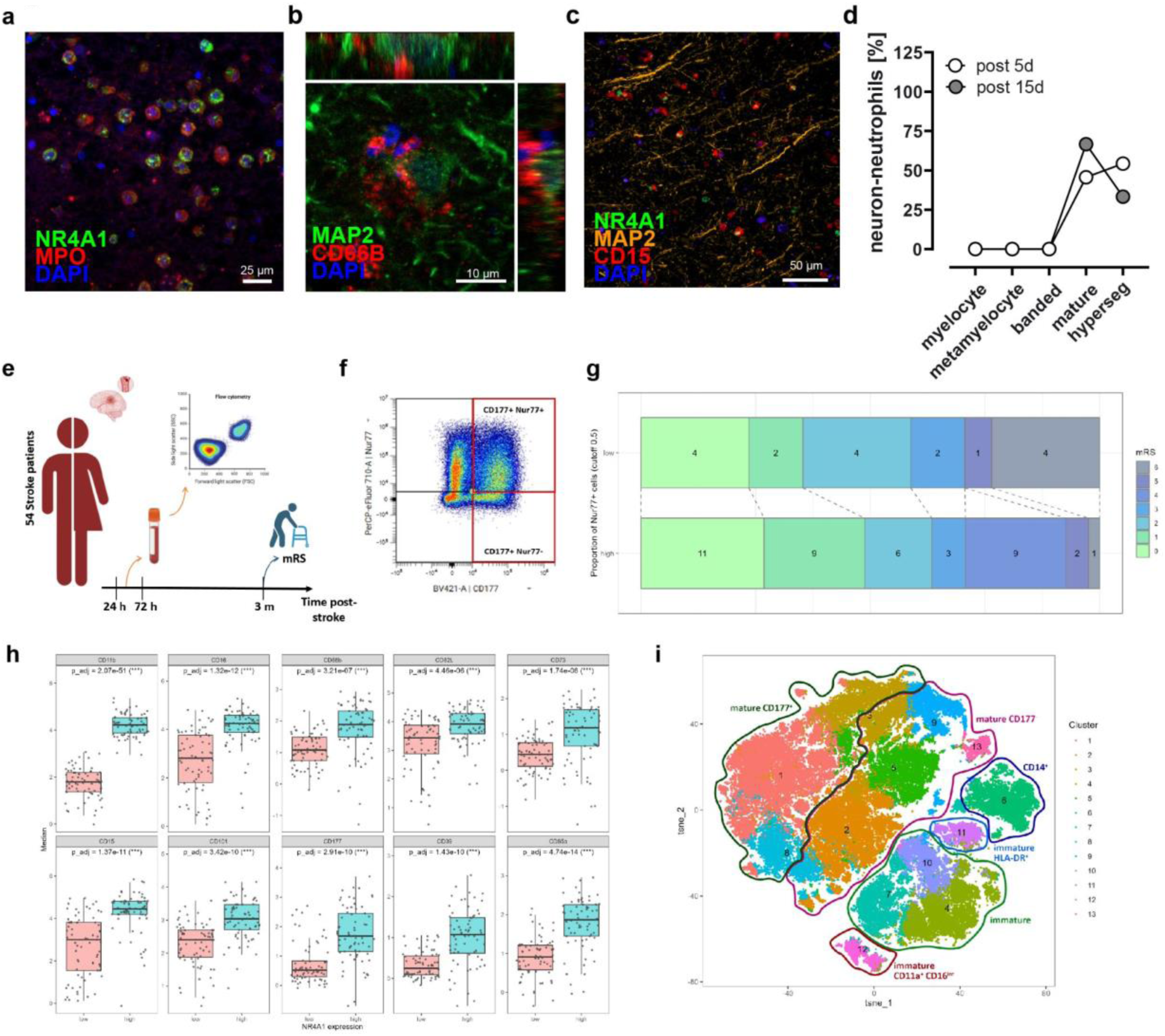
Association between human NR4A1^high^ neutrophils and stroke outcome reveals a distinct NR4A1^high^ phenotype. **a,** Immunofluorescence photographs depicting NR4A1-expressing MPO^+^-neutrophils. in close contact to neurons and neutrophil-specific Ly6b.2-signal within the damaged neurons. **b,** Immunofluorescence image showing a CD66B^+^-neutrophil in close contact to a MAP2^+^-neuron. **c,** MACSima-image showing NR4A1-expressing CD16^+^-neutrophils in close contact to neurons and dendrites. **d,** Quantification and characterization of neutrophils attached to neurons depending on maturation grade within the lesioned brain tissue of a stroke patient five days after symptom onset. **e,** Study design. A cohort of 54 stroke patients was enrolled. Neutrophils were analyzed by spectral flow cytometry between 54 and 72 hours after stroke onset or last known well. Functional outcome was assessed three months post-stroke using mRS. **f,** Flow cytometry gating of human neutrophils was based on CD177 and NR4A1 expression. **g,** Distribution of mRS based on the proportion of NR4A1^high^CD177^+^ neutrophils. **h,** Expression levels (MFI) of neutrophil markers in NR4A1^high^ and NR4A1^low^ neutrophils. The full marker panel is available in Suppl. Fig. 5. **i,** t-SNE representations obtained from human neutrophils. Clusters identified by the unsupervised PhenoGraph algorithm are indicated by colors/numbers. Encirclements depict overarching neutrophil subsets as identified by single marker expressions and/or manual gating.

## Discussion

In this study, we demonstrate that the nuclear receptor NR4A1 plays a critical role in the pathophysiology of ischemic stroke. In response to ischemia, NR4A1 is induced in both mouse and human brains, and its absence in rodents leads to aggravated functional deficits and increased infarct size after experimental stroke. Conversely, ligand-mediated activation of NR4A1 reduced brain inflammation and neuronal damage, establishing NR4A1 as a promising therapeutic target for ischemic stroke. Based on prior evidence from other CNS diseases (Liebmann et al., 2018; Liu et al., 2019), we initially hypothesized that T cells mediate the protective effects of NR4A1 in stroke. Bone-marrow chimera experiments confirmed that NR4A1-dependent effects arise from hematogenous immune cells. However, T cells were not responsible. Instead, neutrophil-specific ablation of *Nr4a1* reproduced the full knock-out phenotype, identifying neutrophils as the principal effector cells. Neutrophils are among the first leukocytes to invade the ischemic brain (Beuker et al., 2021) and contribute to a broad spectrum of immune-mediated diseases. Our findings support two key conclusions: (1) neutrophils are major drivers of immune-associated brain injury following stroke, and (2) NR4A1 acts as a central regulator of neutrophil activation and effector functions, thereby limiting sterile inflammation and improving recovery.

Importantly, several potential mechanisms could be excluded. We found no evidence for NR4A1 effects through enhanced vascular damage due to neutrophil-driven thrombosis, altered neutrophil survival in the CNS, impaired microglial phagocytosis, or increased expression of neutrophil-attracting chemokines. Instead, our functional assays combined with single-cell transcriptomics indicate that NR4A1 functions as a neutrophil-intrinsic brake that limits activation and neurotoxic potential. CNS neutrophils from *Nr4a1*-deficient mice exhibited increased migratory capacity and strong upregulation of proinflammatory programs, including NETosis, ROS production, cytokine signaling, and tissue-destructive pathways. These changes were absent in bone marrow and blood, suggesting that NR4A1 exerts its regulatory function specifically during effector differentiation in the acute sterile inflammatory milieu of stroke. It is conceivable that NR4A1, induced upon stimulation, acts to prevent overshooting neutrophil responses in such settings. Beyond transcriptional and functional alterations, we also identified direct interactions between neutrophils and neurons in ischemic brain tissue from both rodents and humans. These contacts were more frequent in *Nr4a1* knock-outs and associated with increased neuronal apoptosis, consistent with a direct neurotoxic effect of neutrophils. While neuronal injury has largely been attributed to soluble mediators such as ROS, MMP-9, and NETs (reviewed in Jickling et al., 2015; Chen et al., 2021; Denorme et al., 2022; Zuz et al., 2024), our imaging revealed adhesive neutrophil-neuron interactions and vesicular transfer of neutrophil markers into neurons. Such interactions were detectable as early as 24 hours after stroke and persisted for at least 49 days. This previously unrecognized mode of direct cellular engagement adds to established indirect mechanisms such as BBB disruption and cytokine release, highlighting an additional pathway through which neutrophils can mediate neuronal injury. From a translational perspective, it is particularly relevant that the infarct-limiting effects of NR4A1 became evident only after 48 hours, a time frame that overlaps with clinically relevant therapeutic windows. Our patient cohort further underscores the clinical significance, as NR4A1 expression in neutrophils during the acute phase correlated with long-term functional outcome. Likewise, pharmacological activation of NR4A1 in mice ameliorated stroke pathology and improved recovery, confirming its therapeutic potential. NR4A1 agonists have previously shown protective effects in other disease contexts, including osteoarthritis (Xiong et al., 2020), Parkinson’s disease (Yan et al., 2020), and influenza infection (Egarnes et al., 2017). Importantly, Cytosporone B improved outcome when administered 3 hours after stroke onset, underscoring its translational relevance. This aligns well with the delayed kinetics of NR4A1 activation, peaking 48–72 hours post-stroke, a period when secondary neurodegeneration and inflammation are prominent drivers of damage. As Cytosporone B can cross the blood–brain barrier or be delivered via alternatively activated immune cells, neutrophil modulation through nuclear receptors emerges as a feasible therapeutic strategy. Although Cytosporone B itself is unlikely to advance clinically, it provides critical proof of concept that targeted modulation of infiltrating neutrophils can confer neuroprotection and improve recovery. This paves the way for the development of clinically applicable NR4A1-directed compounds.

In summary, our findings identify NR4A1 as a central regulator of neutrophil-mediated injury in ischemic stroke. By restraining proinflammatory effector programs and limiting direct neurotoxic interactions, NR4A1 protects the brain in both mice and humans. Pharmacological modulation of NR4A1 thus represents a promising translational strategy to mitigate immune-driven secondary damage and enhance functional recovery after stroke.

## Material and Methods

### Ethical regulations

Animal procedures were performed in accordance with ARRIVE guidelines and local animal welfare regulations. Experimental protocols were approved by the local governmental authorities (Landesamt für Natur, Umwelt und Verbraucherschutz, NRW, Germany).

### Animals

Adult male and female mice (12–18 weeks of age) with a C57BL/6 background were used in all experiments. Mice lacking *Nr4a1* (Lee et al., 1995, B6;129S2-*^Nr4a1tm1Jmi^*/J) were purchased from Jackson Labs. Male B6.SJL-*Ptprc*^a^*Pepc*^b^/BoyJ (i.e. CD45.1 congenic C57BL/6) mice were used to generate traceable bone marrow chimeras. *Nr4a1*^flox^ mice were kindly provided by Pierre Chambon, Institut de Genetique et de Biologie Moleculaire et Cellulaire, Illkirch-Graffenstaden, France. Mature neutrophil-specific *Nr4a1*-deficient mice were generated by crossing *Ly6g*-Cre heterozygous mice (Ly6g^tm2621(cre)Arte^) with *Nr4a1*^fl/fl^-mice generating either *Ly6g*Cre^+/−^/*Nr4a1*^fl/fl^ or *Ly6g*Cre^−/−^/*Nr4a1*^fl/fl^ animals as controls. Tracking of hematogenous wild type immune cells in mixed-bone marrow mice was done by using C57BL/6-Tg(CAG-EGFP)131Osb/LeySopJ Gfp-reporter mice. Mice were housed under standard laboratory conditions with a 12:12 h light/dark cycle, access to food and water ad libitum in ambient temperature between 20–24 °C and humidity between 45-65%.

### Middle cerebral artery occlusion in mice

Middle cerebral artery occlusion (MCAO) was induced in anaesthetised mice (1-1.5% isoflurane, N_2_O and O_2_) while maintaining a body temperature of 37°C throughout the procedure. After midline neck incision, the left common artery and carotid bifurcation were exposed, and the external and proximal left common arteries were subsequently ligated. Retrograde perfusion of the left common carotid artery was temporarily interrupted using a microvascular clip. The common carotid artery was then incised with micro dissecting scissors, and a silicone-coated 8-0 nylon monofilament (701956PK5Re or 702056PK5Re, Doccol Corporation, Sharon, MA) was advanced to the middle cerebral artery junction. After 30 or 45 minutes of MCA occlusion, the filament was withdrawn to allow MCA reperfusion.

### Middle cerebral artery occlusion in rats

Middle cerebral artery occlusion (MCAO) was initiated in rats under anesthesia (ketamine hydrochloride (100 mg/kg body weight) and xylazine hydrochloride (8 mg/kg body weight, i.p.), maintaining a constant body temperature of 37°C. After a midline neck incision, the left common carotid artery (CCA) and carotid bifurcation were exposed, and the proximal left common and external carotid arteries were ligated. Retrograde perfusion of the left CCA was temporarily interrupted by a microvascular clip (FE691; Aesculap), subsequently the common carotid artery was incised, and the occluder (made from a 2.5 cm long 4.0 nylon suture (Ethilon, Tilburg, the Netherlands) rounded to a diameter of 0.2-0.3 mm using a soldering iron) was advanced to the middle cerebral artery. After 45 minutes of vessel occlusion verified by laser Doppler (Periflux 5001; Perimed), the filament was withdrawn for reperfusion of the MCA. After the procedure, the wound was sutured and treated with xylocain gel (2%).

### Photothrombotic stroke

Photothrombotic stroke was induced under general anesthesia with ketamine hydrochloride and xylazine hydrochloride. Body temperature was maintained constant at 37 °C throughout the procedure. After skull exposure, a laser beam (4 mm diameter, Cobolt Jive TM 75 Laser, 561 nm, Cobolt AB Sweden) was positioned 2 mm to the right of the bregma. Following injection of Bengal Rose (0.1 ml, 10 mg/ml i.p.) the skull was illuminated for 20 minutes for occlusion of cortical vessels.

### Functional testing

For foot fault measurements, animals were individually placed on an elevated 10 mm square wire mesh (total mesh area 40 cm x 40 cm) and then videotaped walking freely for 2 minutes. The number of foot faults and total steps were counted and the percentage of faults was calculated as number of foot faults on the affected limb / number of total steps on the affected limb*100.

### Flow cytometry

Mouse brains were dissected after cardiac perfusion with 20 ml PBS, digested (PBS/ Collagenase, 30 minutes, 37 °C), passed through a cell strainer and washed with PBS/1% FCS. Peripheral immune cells were obtained from cerebral cell suspensions after density centrifugation (2800 x g, 20 minutes in 37 % Percoll). Cells were then washed and resuspended in PBS/2mM EDTA/2% FCS. To obtain splenic single cell suspensions, spleens were passed through cell strainers together with red blood cell lysis (ACK) buffer, centrifuged and resuspended in PBS supplemented with 2% FCS and 2 mM EDTA. For antibodies used see supplementary tables. Dead cells were identified with Fixable Viability Dye eFluor 780 (eBioscience; Thermo Fisher Scientific) or Zombie Green (Cat. No. 423112, 1:500, Biolergend). After staining, cells were washed twice and resuspended in PBS with 2% FCS. Cells were acquired on a Gallios flow cytometer (Beckman Coulter) or FASC Symphony (BD Biosciences). Data were analyzed using Kaluza (Beckmann Coulter) or FlowJo software v10.6.1 (BD Biosciences). Cell concentrations were manually counted in a Fuchs-Rosenthal counting chamber. For spectral flow cytometry cells were stained with fluorescent-labelled antibodies (supplementary table 1) in staining Buffer (4 °C, 20 min, BioLegend). Flow cytometry was performed using the Cytek® Aurora and data was analyzed using FlowJo (Beckton Dickinson) and R Software.

### Bone marrow chimeras

Bone marrow chimeric mice were generated by transplantation of 0.5 x 10^7^ congenic bone marrow cells from either wild-type (C57Bl/6J, CD45.2) or *Nr4a1*-deficient mice (CD45.2) into the coccygeal vein of sublethally irradiated (7.5 Gy) congenic wild type recipients (C57Bl/6J, CD45.1). Successful chimerism was validated after 8 weeks of reconstitution by flow cytometric analysis for allelic CD45 variants in blood samples collected from the tail (CD45.1 vs. CD45.2). Animals with >88% CD45.2+ and <12% CD45.1^+^-leukocytes were used for subsequent experiments.

### Adoptive T cell transfer

For the preparation of splenic lymphocytes, two wild-type and two *Nr4a1*-deficient mice (aged 12-16 weeks) were sacrificed, spleens separated and squeezed through a cell strainer (70 µm mesh). Cells were then washed by centrifugation (300 rpm, 5 min), resuspended in MACS buffer (PBS with 1% FCS, 0.4% EDTA) and again squeezed through a cell strainer (40 µm mesh). Cells were then washed in MACS buffer and T cells were MACS sorted from the suspension using a Pan-T cell isolation kit (Miltenyi Biotec, Bergisch Gladbach, Germany). Subsequently, either sorted wild type or *Nr4a1*-deficient T cells (5×10^6^) were transferred intravenously into *Rag-1*-deficient mice (lacking T and B cells). Experimental cerebral ischemia was induced by MCAO 24 hours after T cell transfer. Sufficient cell transfer was confirmed 24 hours after MCAO by flow cytometric analysis of CD3^+^-T cells in the blood.

### Cytosporone B treatment

Cytosporone B or the corresponding placebo (16.5% DMSO in PBS) was applied to wildtype mice (13 mg/kg body weight, i.p.; Dai., et al., 2014) 3h after induction of reperfusion. *Nr4a1*-deficient mice received either Cytosporone B or the solvent 3h after reperfusion.

### Bicentric study

The study was performed in 2023 in two test centers (Münster UKM, centre 1; Essen UKE, centre 2). Infarct size on day 3 after MCAO was defined as the primary endpoint. Group-assignment (placebo versus active drug) was performed using a random generator (GraphPad Prism - assign subjects to groups). Both active and placebo units were blindly coded and provided by the Münster site. Subsequent infarct detection and functional testing were performed by blinded personnel. Unblinding and categorization was performed after data collection in Münster. The study was pre-registered on 3^rd^ of January 2023 at https://archive.org/details/osf-registrations-865p2-v1. Prior to the study, inclusion and exclusion criteria were defined. Mice were included if the duration of the MCAO procedure was ≤ 60 minutes, sufficient occlusion of the MCA for 30 minutes and successful treatment application. Exclusion criteria were MCAO duration > 60 minutes, insufficient occlusion of the middle cerebral artery, intraoperative bleeding, circling after MCAO, weight loss of ≥ 20% and persistent abnormal posture.

### Tissue collection and processing for histology

Six hours, 24h or 72h after MCAO, mice were perfused through the left ventricle with 30ml cold phosphate buffered saline (PBS) for FACS and mRNA-analyses or for 5 minutes with PBS followed by 4% paraformaldehyde solution for 10 minutes for histology tissue preparation. Mice were held under deep xylazine/ketamine anesthesia during perfusion. Brains were removed and transferred to PBS (FACS) or frozen on dry ice (mRNA analyses). For histological staining, brains were fixed in 4% paraformaldehyde overnight, immersed in 20% sucrose for three days, embedded in TissueTek® frozen and stored at −80°C until further use.

### Infarct volume assessment

Infarct volumes were obtained by collecting 10 µm coronal cryosections in intervals of 300 µm intervals starting at the rostral border of the infarction. Subsequently, slices were stained with 0.5% toluidine blue and dried in graded ethanol (1 min each) at concentrations of 50%, 80%, 96% and 100%. Digitized images were then measured, using ImageJ software, by an investigator blinded to genotype or treatment. Infarct volume was calculated by multiplying infarct area size by 300 µm. Edema compensation was applied using the formula (area of contralateral hemisphere / area of ipsilateral hemisphere)*infarct area.

### Immunohistochemistry

Mounted coronal mouse cryosections were incubated in Blocking Reagent (Roche Diagnostics) for 15 min to prevent nonspecific protein binding. We used the following primary antibodies for murine sections: rabbit-anti-4 Hydroxynonenal (4HNE, 1:100, Abcam, ab46545), rabbit-anti-Cxcl1/Gro-alpha (1:100, Abcam, ab86436), hamster-anti-CD3 (1:50, BDBioscience, 550277), mouse-anti-CEACAM8 (CD66B, 1:100, Invitrogen, MA1-26144), rat-anti-F4/80 (1:500, clone CI:A3-1, Serotec, MCA497G), rabbit-anti-Fibrinogen (1:100, Abcam, ab2413), mouse-anti-GFAP (1:500, clone G-A-5, Millipore, G3893), goat-anti-IBA1 (1:50, Abcam, ab5076), rabbit-anti-Laminin (1:100, Abcam, ab11575), rat-anti-Ly6b.2 (1:100, clone 7/4, BioRad, MCA771G), chicken-anti-MAP2 (1:500, Abcam, ab5392), rabbit-anti-NeuN (1:150, clone 27-4, Millipore, MABN140), rabbit-anti-NGFI-B (NUR77, NR4A1, 1:100, Novus Biologicals, NB100-56745), rabbit-anti-NR4A1 (1:100, Invitrogen, PA5-34033) and mouse-anti-Thrombospondin (1:100, Invitrogen, MA5-13398). Subsequently, brain slices were incubated with the appropriate Alexa Fluor secondary antibody for antigen visualization (1:100, 45 min, room temperature). Cellular nuclei were counterstained using a fluorescent preserving mounting medium containing 4’,6-diamidino-2-phenylindole (DAPI, Life, 00-4952-52). Apoptotic cells were stained by terminal deoxynucleotidyl transferase dUTP nick-end labeling (Click-iT Plus TUNEL Assay, Thermofisher Scientific, C10617).

### Real-Time Polymerase Chain Reaction

Primers used for expression studies were purchased from Qiagen (QuantiTect Assays, Hilden. Germany): *Ccl2* (QT00167832), *Cxcl1* (QT00115647), *Il6* (QT00098875), *Nr4a1* (QT00101017), *Pgk1* (QT00306558), *Ptgs2* (QT00165347), *Tnf* (QT00250999) and the mouse RT^2^-Kit GeneGlobeID-PAMM-181Z by Qiagen. Complementary DNA (cDNA) concentrations were measured by semi-quantitative real-time polymerase chain reactions (RT-PCR) using the BioRad CFX384 RT-PCR-System (Hercules, CA; USA) and SYBR-green fluorescence. Standard LightCycler conditions were used for the expression arrays, and all measurements were done in duplicates. Gene expression was related to the individual expression of phosphoglycerate-kinase (*Pgk1*) as endogenous control. Expression analysis was performed using expression software tool REST-MCS V2 (Pfaffl et al., 2002). In cases when cDNA samples did not exceed the set RT-PCR threshold after 40 cycles (not determined), the respective ΔCT was equated with 40 for the further analysis. RT²-data was analyzed using the Qiagen GeneGlobe analyzer (https://geneglobe.qiagen.com/us/analyze). Stored cDNA-libraries from previous investigations (12h, 18h, 24h, 36h and 168h) have been used for examination of *Nr4a1*-expression after MCAO.

### Single-cell RNA sequencing

All single-cell RNA samples were processed using CellRanger 9.0.0 (Cheng et al., 2017) and the murine reference GRCm39-2024-A for mRNA gene expression data, and a custom reference for custom antibody expression. For each scRNA sample, a matching SingleCellExperiment object was created, and combined into one SCE object. Only genes with an average count above the threshold 0.00001 and expression in at least 5 cells were kept for further analysis. Mitochondrial reads, ERCC spike-in and total reads per were used to flag and remove outliers with SCE’s isOutlier function (Amezquita et al,. 2020), with a threshold of 5% for mitochondrial reads. Doublet cells were filtered using HTO and barcode information, and fastMNN was used for batch correction using d=50 corrected PCs and k=20 MNN neighbours per batch. The data was clustered using the Louvain algorithm, and cell annotation was conducted based on the R/Bioconductor package SingleR (R core team, 2024), with the murine ImmGen reference data set. The annotated scRNA data was subsequently filtered with R v4.3.3 to contain only neutrophil cells, and the remaining cells were reclustered with Seurat v4.4.0 (Satija et al., 2015) and the Louvain algorithm at a resolution of 0.2, and visualized as two-dimensional UMAP. Basic Seurat functions were employed to depict chosen marker genes of interest, to plot average expression scores for gene signatures, and to identify differentially expressed genes (DEG) with the FindMarker function between the main conditions KO and control, using Bonferroni correction and an adjusted p-value cutoff of 0.05. A separate clustering was created for all CNS cells, using Seurat and the Louvain algorithm with a resolution of 0.2 as in the full data set. Relative numbers of KO and control cells per cluster were exported and visualized as barplots with ggplot2 (Wickham et al., 2017) for all organ subpopulations. Focussing on the CNS subset, native Seurat graphic functionality was used to plot a DEG heatmap for clusters and conditions based on the respective top 20 genes per subgroup, and a volcano plot was created with ggplot2 v3.5.2 to show DEGs between KO and control and indicate chosen genes of interest. Enrichment analyses were conducted with clusterProfiler 4.7.1 based on the upregulated genes in the KO condition in comparison to control, and the GO BP database. Significant GO BP terms with an adjusted p-value < 0.05 were grouped into functional categories and displayed as dotplots with ggplot2. Furthermore, enrichment scores for a custom neutrophil signature database were created with the R packages ESCAPE (Borcherding et al., 2021) and UCell (Andreatta et al., 2021) for each cell, and groupwise tests for differences were conducted using ANOVA and Tukey tests between conditions (KO vs control) or clusters (cluster 0 vs remaining clusters; cluster 1 vs remaining clusters). Sequencing was performed by the Core Facility Genomics of the Medical Faculty of the University of Münster.

### ATAC-seq sample and library preparation

Bone marrow (BM) cells were obtained from mice mortaring hip bones, femurs, tibiae and humeri with DPBS. To obtain single-cell suspensions, cells were filtering through a cell strainer of 40 μm diameter. Cell suspensions were incubated for 10 min with ice-cold ACK buffer (0.15 M NH_4_Cl, 10 M KHCO_3_, 0.1 mM EDTA, pH 7.3) for red-blood cell lysis and were washed in DPBS. Labeling of single-cell suspensions was performed on ice in DPBS supplemented with 1% FBS. Cells were first incubated for 10 min in DPBS supplemented with 1% FBS and purified Cd16/32 antibody to block Fc binding. Next, cells were stained against c-Kit (#25-1171-82, ThermoFisher), Cd115 (#135506, BioLegend), Ly6g (#560603, BD Horizon), and CD182 (#149306, Biolegend) for 10 min on ice in the dark. Discrimination of dead cells was performed by addition of 7-aminoactinomycin D (7-AAD; BioLegend cat#:420404) viability stain solution 10 min before measurement. ATAC-seq was performed on FACS (Aria III, BD Biosciences) sorted cKit^neg^CD11b^pos^Ly6g^pos^CD182^low^ and cKit^neg^CD11b^pos^Ly6g^pos^CD182^pos^ primary BM neutrophils as previously described (Corces et al., 2017), with the exception that KAPA HiFi HotStart ReadyMix (Roche, cat#:7958927001) was used as polymerase. Quality was assessed on a High Sensitivity DNA chip (Agilent Technologies). Quantity was measured using the NEBNext Library Quant Kit for Illumina (New England Biolabs, cat#:E7630S). Libraries were sequenced paired-end on NextSeq2000 (P2, 200 cycles). Sequencing was performed by the Core Facility Genomics of the Medical Faculty of the University of Münster.

### Electron Microscopy

Mice were transcardially perfused with 2% (v/v) formaldehyde and 2.5% (v/v) glutaraldehyde in 100 mM cacodylate buffer (pH 7.4). Bone marrow cells were extracted from the tubular bones using a cannula and PBS, passed through a 70-µm cell strainer, centrifuged (300×g, 10 min, 4 °C), and washed twice in PBS. Subsequently, cells were post-fixed in 0.5% (v/v) osmium tetroxide and 1% (w/v) potassium hexacyanoferrate (III) in 0.1 M cacodylate buffer (2 h, 4 °C), followed by washing with distilled water. After dehydration in ascending ethanol concentrations (30–100%), cells were incubated twice in propylene oxide (15 min each) and embedded in Epon using Beem capsules. Ultrathin sections were collected on copper grids and negatively stained with 2% uranyl acetate for 10 min. Electron micrographs were acquired at 60 kV with a Philips EM-410 electron microscope using imaging plates (Ditabis, Pforzheim, Germany).

### Sample size calculation

Sample sizes were calculated with sample size calculator (http://www.stat.ubc.ca). A priori sample size calculations were set to achieve 80% power to detect a relevant treatment effect of 20% at α=0.05.

### Human samples

#### Ethics statement

The study was conducted in accordance with the Declaration of Helsinki. With the approval of the Ethics Committee of University of Münster for the NeutroStroke study (registration no. 2022-077-f-S), informed consent was obtained from healthy donors and patients or their legal representatives.

#### Histology

For histological analyses, brain autopsy material from five patients (three women and two men, with a mean age of 77.2 years and an age range of 62–93 years) who died from acute ischemic stroke between two and fifteen days (median five days) after stroke onset at the University Hospital of Münster was analysed. Brain samples were derived from brain regions within the vascular territory of the middle cerebral artery (cortex, striatum and internal capsule). Prior to postfixation an approximately 2 cm thick coronal brain slice was taken at MNI coordinate y = 0 mm, corresponding to the midline between the anterior and posterior poles of the brain and stored at −80 °C.. The anterior and posterior brain sections were then fixed in paraformaldehyde (PFA) for 14 days until preparation of tissue slides. Frozen sections were fixed in PFA (4 °C overnight) and dehydrated for 6 weeks in 30% sucrose. The sucrose solution was replenished regularly. Subsequently, tissue was cut with a cryostat (10 µm) and frozen at −20 °C until further use.

#### Study design, patients and participants

Adult patients were recruited with either imaging-confirmed ischemic stroke or transient ischemic attack (TIA). Imaging confirmation was obtained via computed tomography (CT) or magnetic resonance imaging (MRI). Patients were excluded if they had severe acute infectious diseases, active malignancies, recent trauma, surgery in the last six months, severe bleeding requiring a blood transfusion in the last three months, pregnancy or were breastfeeding. Blood samples were collected in the morning between 24 and 72 hours after the onset of stroke symptoms or the time they were last known to be well. Neutrophils from 54 patients were analyzed by spectral flow cytometry. 12 patients were excluded from the study in a retrospective manner due to a change in diagnosis or delayed notification of exclusion criteria and 12 patients were excluded retrospectively due to low sample quality in the flow cytometry.

#### Clinical assessment

All patients included in the study underwent regular clinical and radiological assessments. Clinical parameters encompassed demographic information, medical history, a thrice-daily assessment of the National Institutes of Health Stroke Scale (NIHSS), and laboratory values such as leukocyte count. The modified Rankin scale (mRS) was assessed via telephone three months after admission.

#### Blood sample preparation for spectral flow cytometry

Directly following the collection of blood samples in EDTA tubes, 100 µL whole blood was processed for spectral flow cytometry analysis. Red blood cells (RBCs) were lysed using 2 mL RBC Lysis Buffer (BioLegend) for 20 min at 4 °C. The lysis process was halted using HANKs buffer (10% HBSS (Gibco), 0.6% BSA Stock Solution (Miltenyi Biotec), 0.06% EDTA 0.5 M (invitrogen) in aqua (sterile, B. Braun), pH-adjusted to 7.4). Subsequent to two washing steps, a viability staining with Zombie NIR (BioLegend) in phosphate buffered saline (PBS, Sigma-Aldrich) was performed for 30 min at 4 °C. Following two additional washing steps, leukocytes were stained with fluorochrome-conjugated antibodies (Table X1) for 30 min at 4 °C. Titration of all antibodies was performed to ensure optimal concentrations. Furthermore, to assess the expression of NR4A1, after another two washing steps the leukocytes were incubated for 10 min in Fixation buffer (True Nuclear Fix Buffer, BioLegend), followed by 30 min of incubation in Permeabilization Buffer (True Nuclear Perm Buffer, BioLegend) containing the fluorochrome-conjugated NR4A1 antibody. Samples were measured on an Aurora spectral flow cytometer (Cytek Biosciences) under daily quality control by SpectroFlo QC Beads (Cytek Biosciences). Longitudinal stability of the measurements was further validated by the repeated analysis of reference samples during the entire study.

#### Preprocessing of flow cytometry data

Flow cytometry data were unmixed using appropriate single reference controls (SRCs) using SpectroFlo™ software (Cytek Biosciences) and further processed using OMIQ software (Dotmatics, www.omiq.ai). Automated data cleaning was conducted using peak extraction and cleaning oriented quality control (PeacoQC) algorithm (1). To exclude remaining doublets and cell aggregates, only events showing linear forward scatter height / area (FSC-H / FSC-A) and side scatter height / area (SSC-H / SSC-A) correlations were included as singlets. Any remaining RBCs were extracted based on the differential side scatter between violet and blue lasers due to hemoglobin absorption. Subsequently, fluorochrome parameters were linearized using arcsinh transformation with a cofactor of 6000. An optimized T-distributed stochastic neighbor embedding (opt-SNE) (2) map was utilized for the identification of neutrophils, along with the expression CD15 and CD66B.

#### Automated gating / tSNE and Phenograph

To investigate complex neutrophil phenotypes in high-dimensional space, a tSNE (3) map was employed to visualize neutrophil subpopulations (iterations: 1000, learning rate: 5000, perplexity: 0.5). Subsequent unsupervised clustering was performed using the PhenoGraph algorithm (4). The number of nearest neighbors (‘k’) was tested up to 700 and we selected k=600, as it introduced a desirable level of overclustering, allowing for the detection of subtle patterns and relationships while maintaining overall cluster stability.

#### Neutrophil counts

Neutrophil counts were determined by multiplying the relative proportion of neutrophils in the flow cytometry data with the leukocyte count obtained from clinical routine diagnostics.

#### Statistics

Unless indicated otherwise, the statistical and computational analysis was performed in RStudio (2024.12.1) running R4.4.3. To assess the relationship between the proportion of CD177^+^NR4A1high neutrophils and the functional outcome measured by mRS, an ordinal logistic regression was performed using R package ‘MASS’ (7.3.65). As independent variables, the proportion of CD177^+^NR4A1^high^ neutrophils, age, sex, initial NIHSS score and neutrophil count were included. Continuous variables were Z-score normalized. A two-tailed t-test was used to compare NR4A1^high^ and NR4A1^low^ neutrophils. The resulting p-values were corrected for multiple testing using the Benjamini-Hochberg method and are visualized as *<0.05, **<0.01, and ***<0.001.

### In vitro studies

#### Cell death analysis

To analyze cell death, HoxB8 cells (5 x 10^5^ cells per sample) were collected at day 0, 3, 4, 5 and 6 of differentiation. Cells were stained with FITC-Annexin V (Biolegend), SYTOXTM Red dead cell stain (Thermo Fisher) and tetramethylrhodamine methyl ester (TMRM, Thermo Fisher). Fluorescence minus one (FMO) controls were used to determine the negative cell population. Samples were analyzed using a CytoFLEX S flow cytometer (Beckman Coulter) and analyzed with FlowJoTM software (BD Biosciences).

#### Flow chamber assays

Induction of adhesion under flow was performed as described before (Pick et al. 2017; Zehrer et al. 2018). Briefly, differentiated HoxB8 cells (7.5 x 10^5^ cells per sample) were perfused through µ-slides VI0.1 (Ibidi) coated with recombinant murine (rm) P-selectin (5 µg/ml, R&D system), rmICAM-1 (3 µg/ml, Stemcell), and rmCXCL-1 (5 µg/ml, Peprotech). Time-lapse images were captured for 9 minutes under constant shear stress (1 dyne/cm²) from 18 different viewpoints using an Axiovert 200M microscope (Zeiss) equipped with a Plan-Apochromat 20 x/0.75 NA objective (Zeiss), an AxioCam HR digital camera, and a temperature-controlled environmental chamber. Quantification of adherent cell numbers after 1, 3, 5, 7, and 9 minutes was performed offline with FIJI software (Schindelin et al. 2012). Cells were considered adherent if stationary for over 15 seconds. Analysis of three-dimensional chemotaxis in collagen gels was analyzed as described before (Salvermoser et al. 2018; Zehrer et al. 2018, J Immunol). The middle channel of Ibidi µ-slides Chemotaxis (Ibidi) was filled with 3 x 10^5^ HoxB8 cells in 1.5 mg/ml rat tail collagen type I (Ibidi) and incubated for 5 min at 37 °C. Cells were exposed to a gradient of rmCXCL-1 (100 ng/ml, Peprotech) and time-lapse videos were recorded every 14 s for 30 min at 37 °C using an Axiovert 200M microscope (Zeiss) equipped with a Plan-Apochromat 10 x/0.3 NA objective (Zeiss). Migration tracks were analyzed manually using FIJI software (Schindelin et al. 2012) and its implemented Manual Tracking plugin (Fabrice Cordeliès, Institute Curie, France). Migration velocity, distance and directionality were calculated using the Chemotaxis Tool (Ibidi).

#### Phagocytosis

For phagocytosis assays 2 x 10^5^ HoxB8 cells were seeded into individual wells of a 96 well plate in Hank’s Balanced Salt Solution (HBSS, Biochrom), supplemented with 20 mM Hepes (Sigma-Aldrich), 0.25% BSA (Sigma-Aldrich), 0.1% glucose (Sigma-Aldrich), 1.2 mM Ca^2+^, 1 mM Mg^2+^ (Sigma-Aldrich) and 10% fetal calf serum (Sigma-Aldrich). 2 x 10^6^ Escherichia coli MG1655 (multiplicity of infection: 10) carrying the plasmid pEB2-E2-Crimson were added per well and incubated with the cells for 30 min at 37 °C. Where indicated, cytochalasin D (10 µg/ml, AppliChem) was added to prevent phagocytosis. Cells were then centrifuged, resuspended in LIVE/DEAD Fixable Violet Dead Cell Stain (Thermo Fisher Scientific) in PBS, incubated for 15 min on ice, and fixed with 2% formaldehyde solution. Samples were analyzed using a CytoFLEX S flow cytometer (Beckman Coulter) and 10^4^ cells were acquired per sample. Subsequent data analysis was performed using FlowJo 10 software (BD Biosciences). Median fluorescence intensity (MFI) values of all single living cells were calculated and results are shown after subtraction of the MFI from the samples incubated with cytochalasin D. pEB2-E2-Crimson was a gift from Philippe Cluzel (Addgene plasmid #10401; Balleza et al. 2018).

#### Metabolic assay

Metabolic activity of HoxB8-immortalised wild type or *Nr4a1*-deficient granulocytes (Gran et al., 2018) was determined using the XFe96 Extracellular Flux Analyzer (Agilent Technologies), as previously described (Liebmann et al., 2018). After five days of differentiation, 200.000 HoxB8-immortalised granulocytes were cultured in XF Base Medium Minimal Dulbecco’s modified Eagle’s medium (Agilent Technologies) containing 10 mM glucose, 2 mM L-glutamine and 1 mM sodium pyruvate (Merck). Oxygen consumption rate (OCR) and extracellular acidification rate (ECAR) were assessed under basal conditions and in response to 2 μM oligomycin, 1 μM FCCP, 100 nM rotenone plus 1 μM antimycin A (Sigma-Aldrich). Cells were pre-treated with or without 600 nM phorbol-12-myristat-13-acetate (PMA, Sigma-Aldrich) for 3h and OCR and ECAR were determined in parallel. OCR and ECAR were analyzed using Wave Desktop software (Agilent).

#### ICAM-1 and fibrinogen binding assay

ICAM-1- and Fibrinogen-binding assays were performed as previously described (Jakob et al. 2013; Lefort et al. 2012). To assess LFA-1-specific ICAM-1 binding, 2×10^5^ murine PMNs murine neutrophils were resuspended in Hanks Balanced Salt Solution (with 10mM HEPES, 1 mM CaCl_2_ and MgCl_2_) and preincubated with a functional blocking anti-Mac-1 antibody (clone M1/70, 10 μg/ml). Neutrophils were stimulated with Cxcl1 (100 ng/ml, 3 min, 37°C, Peprotech), or left untreated in presence of anti-ICAM-1/Fc (20 μg/ml, clone B3.3) and APC-conjugated anti-human IgG1 (Southern Biotechnology). Afterwards, neutrophils were fixed on ice (7.4% formaldehyde) and stained with Ly6b.2 antibody (Bio-Rad). Mean fluorescence intensity of LFA-1–specific binding of ICAM-1/Fc was calculated on Ly6b.2^+^-cells. To investigate Mac-1 binding to Fibrinogen, isolated murine neutrophils were incubated in 0.9% NaCl (with 0.1% glucose, 0.25% BSA, 2 mM HEPES) with 150 μg/mL Alexa 647-conjugated fibrinogen (Invitrogen) and stimulated with Cxcl1 (100 ng/ml, 10 min, 37°C, Peprotech), or left unstimulated. Neutrophils were then stained with Ly6b.2 (Bio-Rad) and the percentage of PMNs positive for fibrinogen binding was calculated by defining a threshold of the fluorescence intensity where 5% of PMNs in the wild type control were considered negative. Analyses of Icam-1- and fibrinogen-binding assay were performed using FlowJo (FlowJo LLC, USA).

## Acknowledgements

We thank Annika Engbers, Thorsten König, Christine Salin, Nina Kreienkamp, Dirk Reinhardt, Heike Hater, Karin Gäher and particularly Maike Hoppen and Birgit Schmeddes for excellent technical assistance. This work was supported by the Interdisciplinary Center for Clinical Research (IZKF) Münster, grant Min3/003/21, and by the Deutsche Forschungsgemeinschaft (DFG; MI 1547/3-1 and FOR 2879/1). LK was supported by grants from the Deutsche Forschungsgemeinschaft (DFG; KL 2199/5-1). This work was further supported by the German Research Foundation collaborative research grant TRR332 (project # 449437943; projects A1 (C.S-R., C.-O.S.), projects A2 (O.S., R.C., M.R.), projects B2 (L.K. and J.M.), projects B3 (H.A., F.R.), projects B5 (J.R.), projects C3 (D.M.-B. and B.W.), projects C6 (D.H. and M.G.), projects C7 (T.V.) and projects Z1 (O.S, D.R.E and K.D.). All sequencing data were generated by the Core Facility Genomics of the University of Muenster.

## Author contributions

Conception and design of the study: JKS, ML, ML, CSR, FR, OS, LK, JM. Providing materials and reagents: TK, CT, TV, SG, JR, PC, FR, BW, DHM, GMzH, MG, AZ, OS, CSR Acquisition and analysis of data: JKS, YS, ML, CW, SR, HA, ALB, ASP, CB, AMY, StH, JR, COS, LK, ASS, MH, MR, JNH, SH, UH, NH, AB, JG, DMB, RC, GMzH, CSR, LK, JM. Drafting manuscript and figures: JKS, ML, LK, JM. Revision and approval of manuscript: all authors.

## Declarations

The authors declare no competing interests. All sequencing information will be made available by uploading to a public repository before publication of the manuscript.

## Supplementary Figures

**Supplementary Figure 1.**
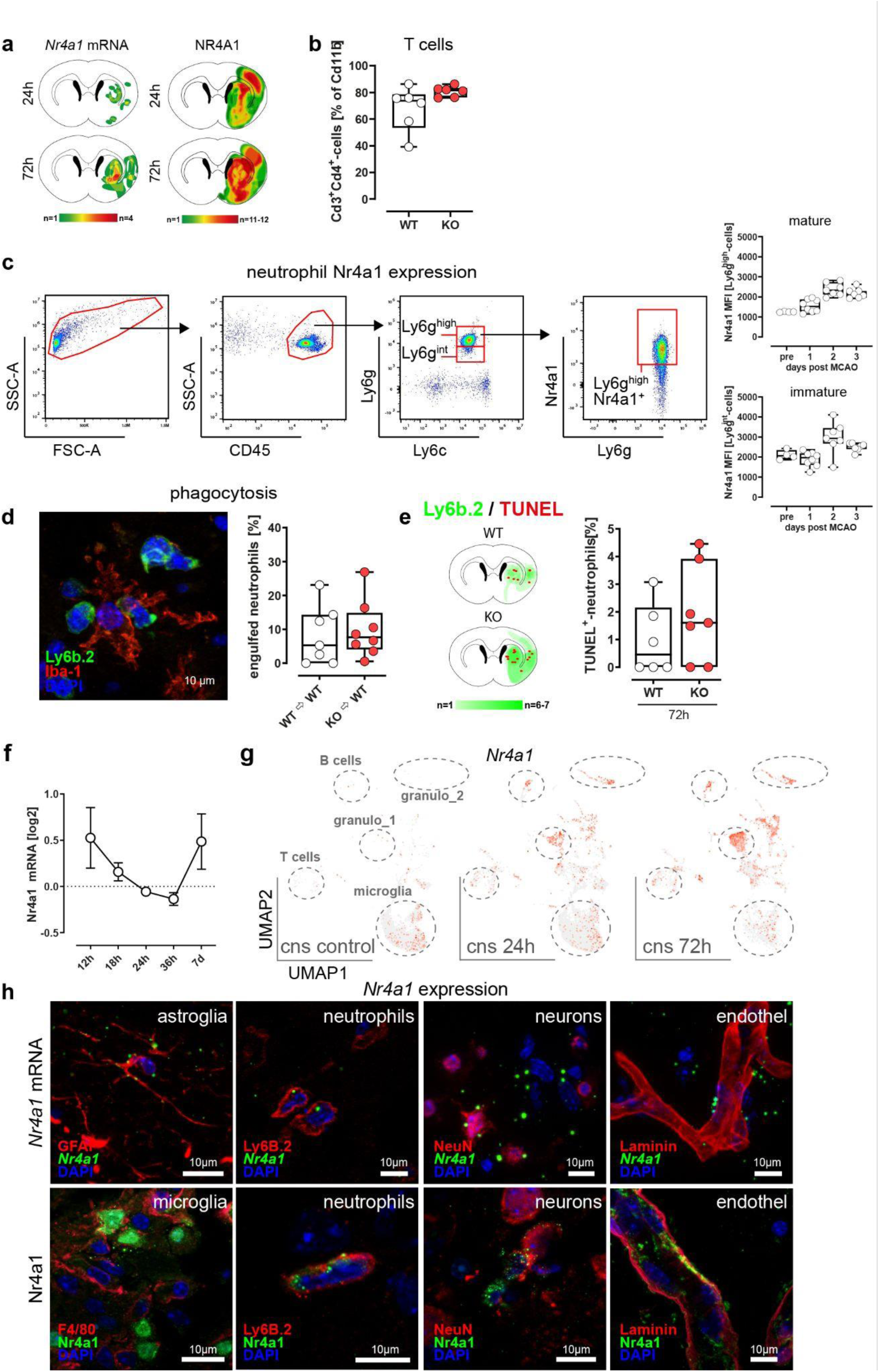
**a,** *Nr4a1*-mRNA and Nr4a1-protein signal heatmaps were generated using RNA-ISH and immunofluorescence-stained coronal sections of wild type mice post 24h (n=4 mRNA, n=12 protein), post 72h (n=4 mRNA, n=11 protein) after experimental occlusion of the MCA. **b,** Quantification of CD3^+^CD4^+^-T cells 72h after MCAO. Median cell counts are shown as box plots. Lines within the boxes denote medians (WT n=6; KO n=6). **c,** Representative gating strategy of NR4A1 expression in neutrophils of wild type mice following MCAO. Strategy applies to the data shown in figure 1i. right panel: NR4A1 expression of immature Ly6g^int^ and mature Ly6g^high^-neutrophils over the course of three days after MCAO. **d,** Quantification of neutrophils either engulfed or phagocytized by microglia within the ischemic hemisphere was performed on cryosections at bregma 0-1 mm 72h post-MCAO (WT►WT n=7; KO►WT n=8). **e,** Quantification and heatmap generation of TUNEL+-apoptotic neutrophils was performed on coronal sections within the ischemic striatum and cortex. Median cell counts are presented as box plots. The lines inside the boxes denote medians (WT n=6; KO n=7). **f,** *Nr4a1*-mRNA expression was assessed in brain lysates over the course of 7 days post MCAO (sham, post 12h, 18h, 24h, 36h and 7d). Each time-point n=4). **g,** *Nr4a1*-feature plot overlaid on the Merged Uniform Manifold Approximation and Projection (UMAP) plot showing cell clusters identified in combined single-cell transcriptome obtained from the brain of sham (cns control) and mice subjected to experimental stroke (24h and 72h post MCAO). Two distinct neutrophil subpopulations emerging after experimental stroke are highlighted. Sequencing raw data are available at GEO GSE189432. **h,** Co-localisation studies of *Nr4a1*-mRNA and protein were performed on mice stroke specimens. Double staining was done with marker for astroglia (Gfap), microglia and monocytes/macrophages (F4/80), neurons (NeuN), blood vessels (Laminin) and neutrophils (Ly6b.2 clone 7/4).

**Supplementary Figure 2.**
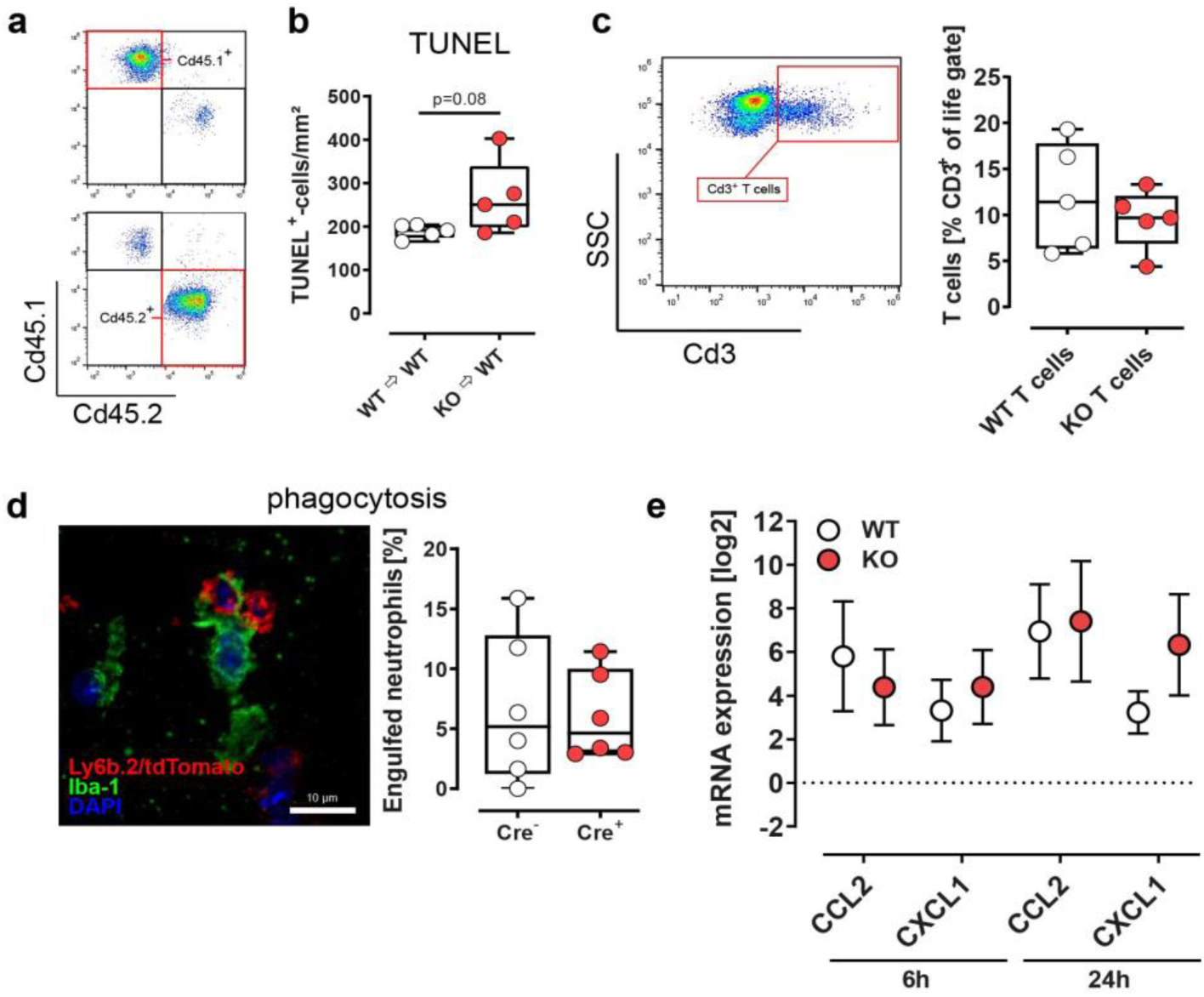
**a,** Representative FACS-plot of the chimerism approach. Cd45.1 congenic wild-type mice (WT) or Cd45.2 *Nr4a1*-deficient (KO) were sublethally irradiated and reconstituted with either congenic wild-type (WT Cd45.2►WT Cd45.1) or *Nr4a1*-deficient (KO Cd45.2►WT Cd45.1) bone-marrow cells (WT Cd45.2►KO; Cd45.1). **b,** Quantification of TUNEL^+^-cells was performed on coronal sections within the ischemic striatum and cortex at 20x magnification. Median cell counts are presented as box plots. The lines inside the boxes denote medians (each group n=5; P=0.08, two-sided *t*-test). **c**, Applied T cells were tracked by FACS analysis of peripheral blood. **d,** Quantification of neutrophils either engulfed or phagocytized by microglia within the ischemic hemisphere was performed on cryosections at bregma 0-1 mm 72h post-MCAO (WT►WT n=7; KO►WT n=8). **e,** Expression of *Ccl2* and *Cxcl1* 6h and 24h after experimental stroke in WT and KO (n=5 each) mice compared to sham operated animals (n=3).

**Supplementary Figure 3.**
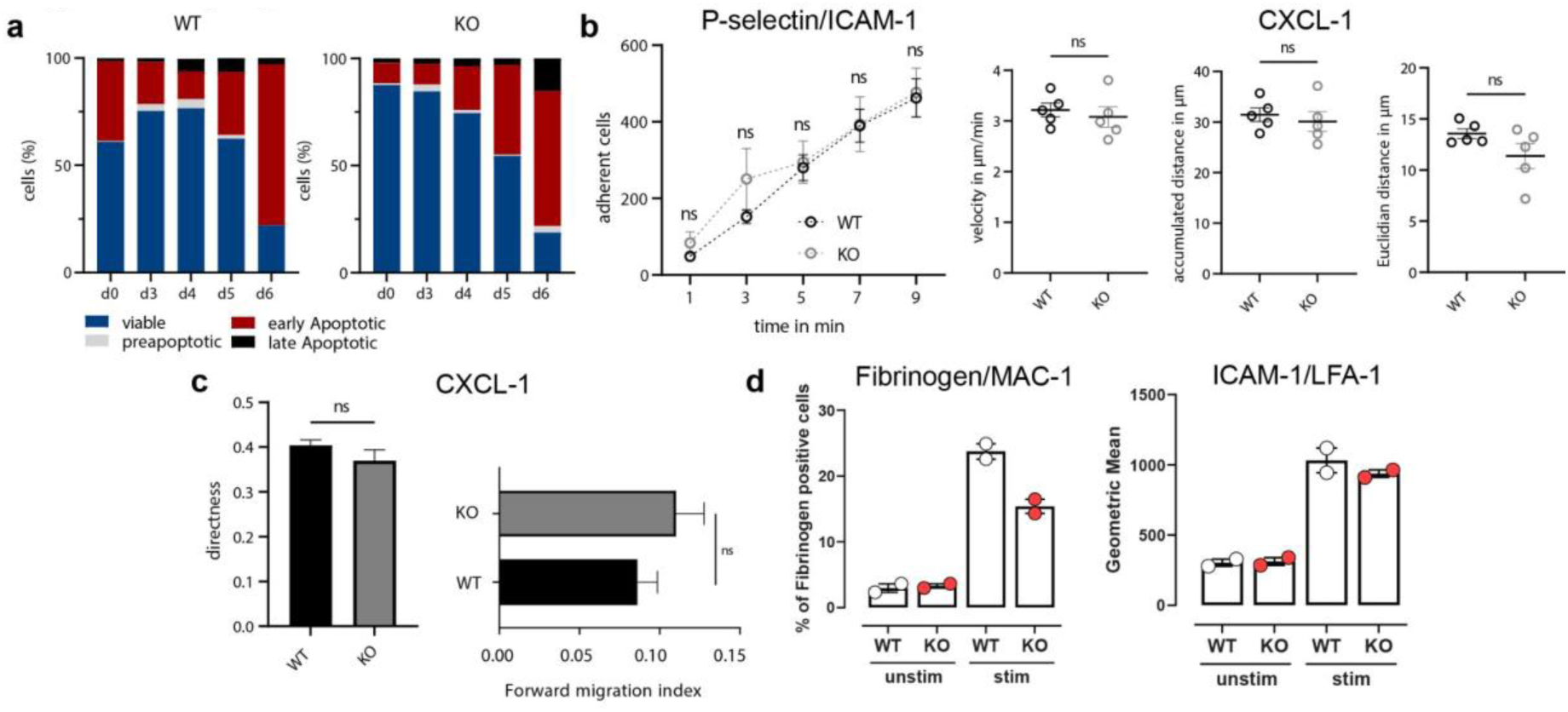
**a,** Cell death analysis was performed in wild type and *Nr4a1*-deficient HoxB8-neutrophils over the time course of 6 days. **b,** Wild-type and *Nr4a1*-deficient HoxB8-neutrophils were compared for adherence (Pecam-1, Icam-1, Cxcl-1) and migration capacity. **c,** Three dimensional chemotaxis assays (Cxcl-1) were performed in wild type and *Nr4a1*-deficient HoxB8-neutrophils. **d,** Icam-1 and Fibrinogen binding assays were performed with sorted bone marrow neutrophils of wild type and *Nr4a1*-deficient mice.

**Supplementary Figure 4.**
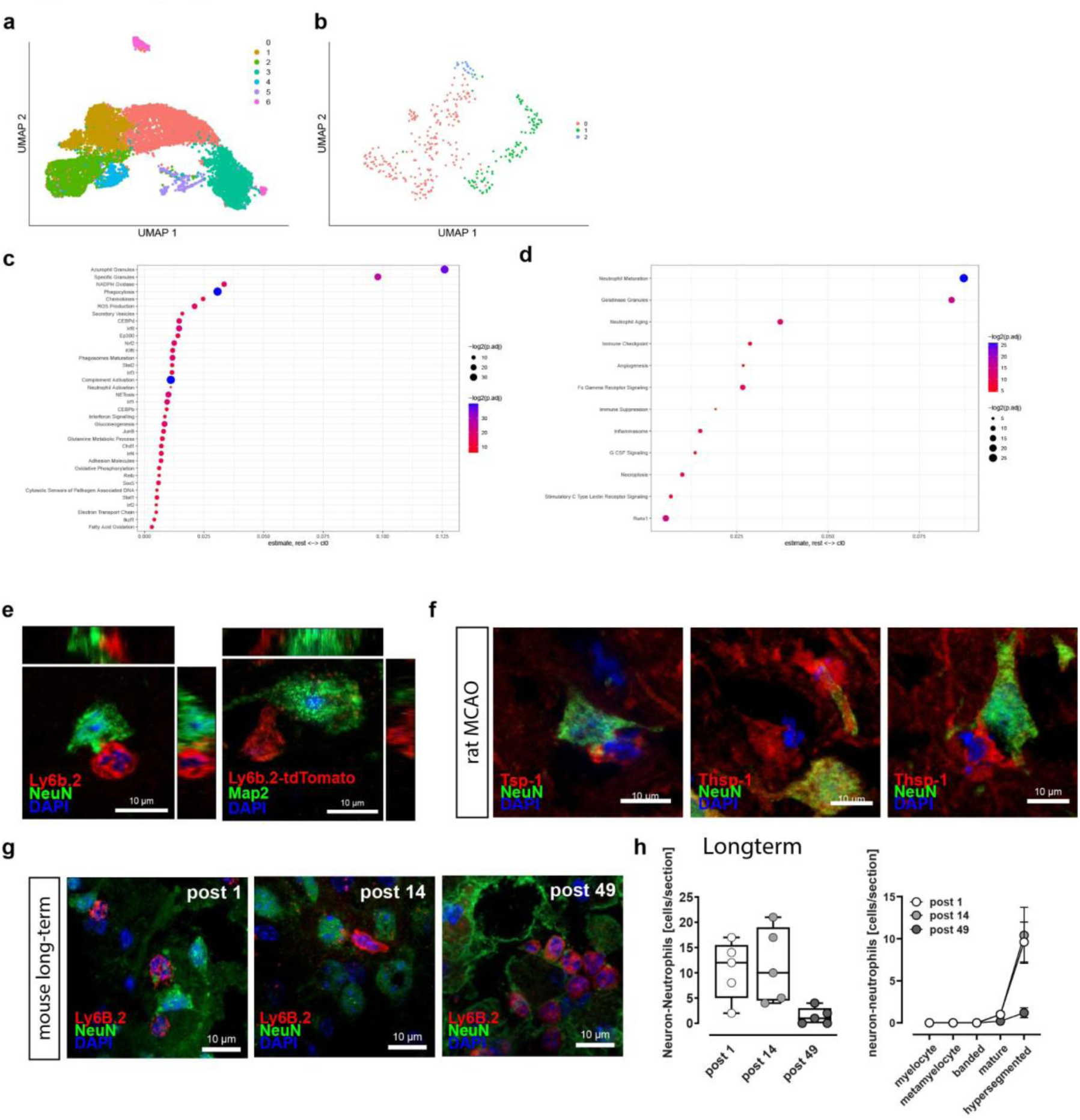
**a,** Single-cell transcriptomics analysis of *Nr4a1*-deficient and wild type mice (n=8). UMAP plot for blood and bone-marrow with Louvain clustering **b,** UMAP plot with Louvain clustering for CNS neutrophil scRNA data **c,** Enrichment analyses of custom neutrophil signatures for CNS clusters 0 and 1. All enrichment scores per cell were based on the ESCAPE/UCell algorithm. ANOVA and Tukey tests (adj.p < 0.05) were applied to test for differences between the respective cluster of choice and the remaining clusters (left: cluster 0 vs clusters 1+2. **d,** cluster 1 vs clusters 0+2). **e,** Image showing Ly6b.2-expressing neutrophil interacting with a neuron within the lesion. **f,** Neutrophil-neuron interaction was confirmed immunohistochemically in rats subjected to MCAO and in **g,** mice subjected to photothrombotic cortical stroke over the time course of 49 days. **h,** Immunohistological neutrophil quantification within the ipsilateral hemisphere post 1d, 14d and 49 days after photothrombotic stroke (each group n=5). Median cell counts are presented as box plots and lines inside the boxes denote medians.

**Supplementary figure 5.**
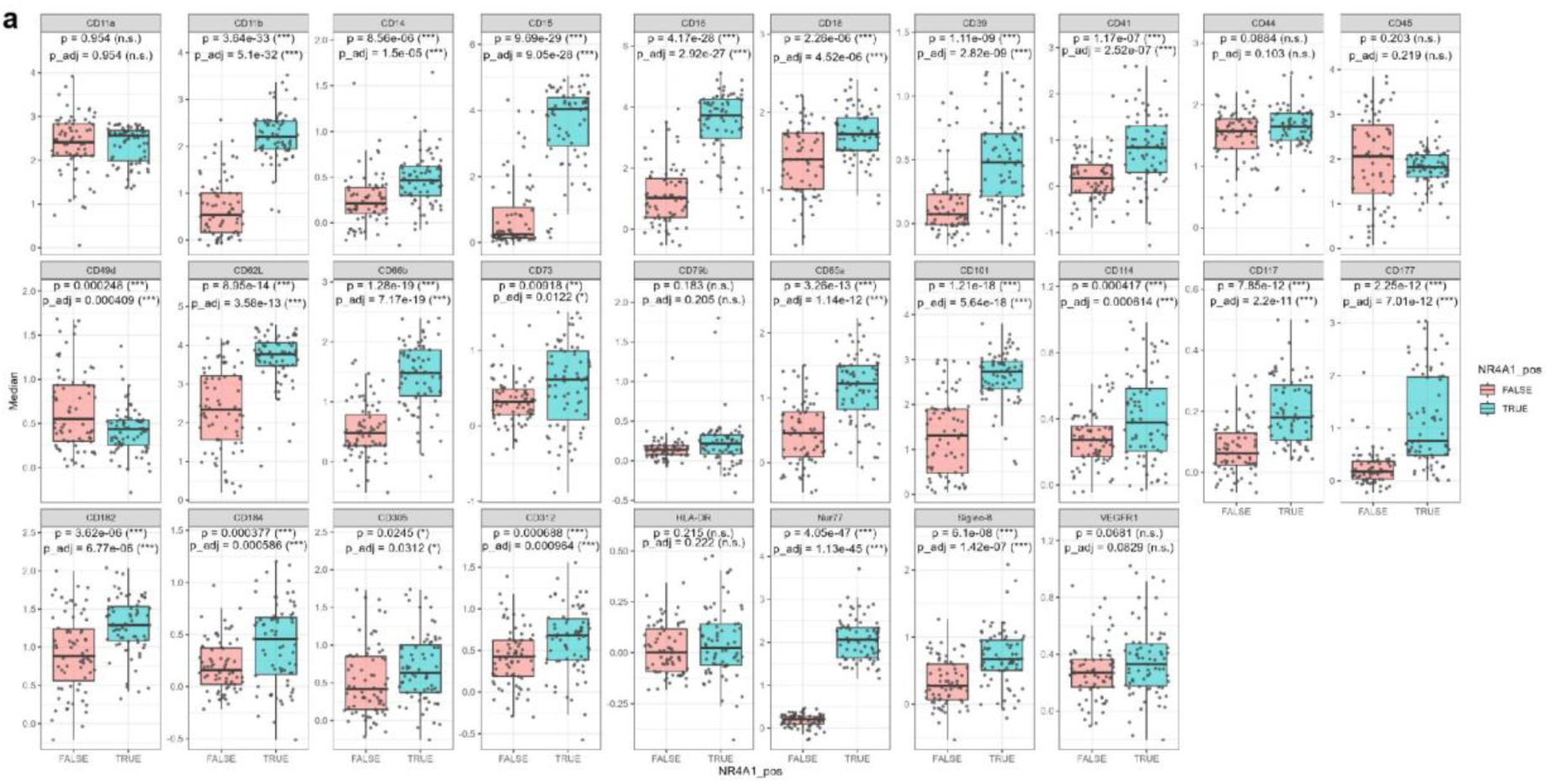
**a**, Full panel of expression levels (MFI) of neutrophil markers in NR4A1^high^ and NR4A1^low^ neutrophils.

**Table 1.** Spectral flow cytometry antibodies.

| Target (clone) | Vendor | Catalogue number | RRID |
| --- | --- | --- | --- |
| CCR2 (475301) | BD Biosciences | Cat# 747967 | AB_2872428 |
| CD101 (Moushi101) | eBioscience | Cat# 17-1011-82 | AB_2815082 |
| CD106 (429) | BD Biosciences | Cat# 745193 | AB_2742789 |
| CD115 (AFS98) | Biolegend | Cat# 135512 | AB_11218983 |
| CD115 (AFS98) | BD Biosciences | Cat# 750948 | AB_2875029 |
| CD117 (2B8) | Biolegend | Cat# 105828 | AB_11042456 |
| CD11b (M1/70) | BD Biosciences | Cat# 749864 | AB_2874105 |
| CD14 (Sa14-2) | Biolegend | Cat# 123332 | AB_2734180 |
| CD16 (S17014E) | Biolegend | Cat# 158012 | AB_2876543 |
| CD16/32 (Ab93) | BD Biosciences | Cat# 751701 | AB_2875686 |
| CD24 (M1/69) | BD Biosciences | Cat# 101836 | AB_2566730 |
| CD34 (700011) | R&D | Cat# FAB6518T | AB_3651676 |
| CD41 (MWReg30) | Biolegend | Cat# 133918 | AB_2563500 |
| CD44 (IM7) | Biolegend | Cat# 103037 | AB_10900641 |
| CD45 (30-F11) | Biolegend | Cat# 103130 | AB_893339 |
| CD49d (9C10) | BD Biosciences | Cat# 741243 | AB_2870794 |
| CD62L (MEL-14) | BD Biosciences | Cat# 746726 | AB_2743990 |
| CD80 (16-10A1) | Biolegend | Cat# 104734 | AB_2563113 |
| CD81 (Eat-2) | Biolegend | Cat# 104912 | AB_2562995 |
| CXCR2 (SA044G4) | Biolegend | Cat# 149304 | AB_2565692 |
| CXCR2 (SA044G4) | Biolegend | Cat# 149314 | AB_2734211 |
| CXCR4 (L276F12) | Biolegend | Cat# 146517 | AB_2687244 |
| CXCR4 (L276F12) | Biolegend | Cat# 146517 | AB_2687244 |
| Ly6C (HK1.4) | Biolegend | Cat# 128041 | AB_2565852 |
| Ly6G (1A8) | BD Biosciences | Cat# 565964 | AB_2739417 |
| MHC II (M5/114) | BD Biosciences | Cat# 750280 | AB_2874471 |
| Sca1 (D7) | BD Biosciences | Cat# 741466 | AB_2870934 |

**Table 2.** Histo antibodies.

| Target (clone) | Vendor | Catalogue number |
| --- | --- | --- |
| 4HNE | Abcam | ab46545 |
| CD66B | Invitrogen | MA1-26144 |
| GFAP (clone G-A-5) | Millipore | G3893 |
| F4/80 (clone Cl:A3-1) | Serotec | MCA497G |
| FIBRINOGEN | Thermofisher | MA5-15902 |
| IBA-1 | Abcam | ab5076 |
| LAMININ | Abcam | ab11575 |
| LY6B.2 (clone 7/4) | BioRad | MCA771G |
| MAP2 | Abcam | ab5392 |
| MPO | Abcam | ab208670 |
| NEUN (clone27-4) | Millipore | MABN140 |

**Table 3.**
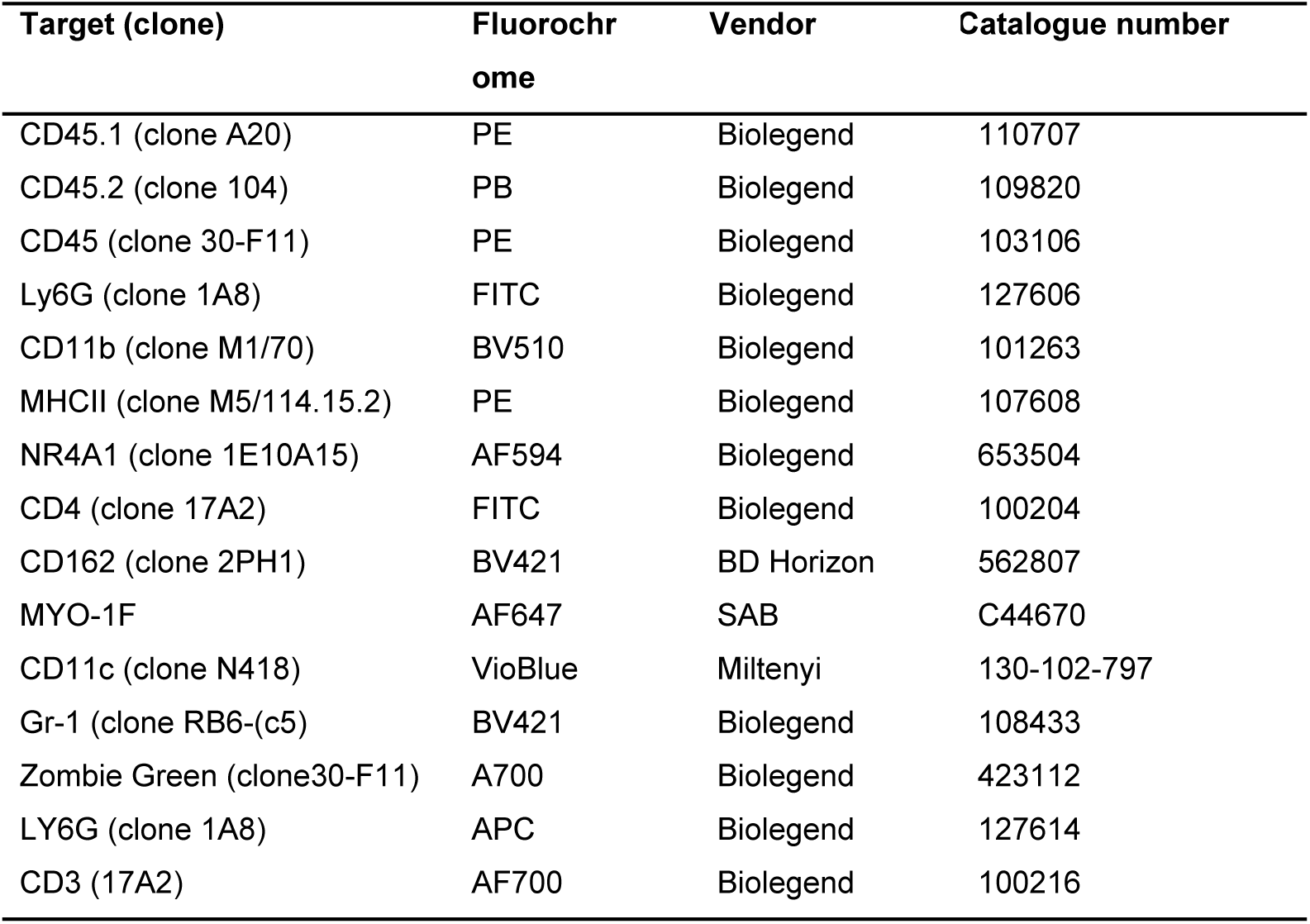
FACS antibodies.

| Target (clone) | Fluorochrome | Vendor | Catalogue number |
| --- | --- | --- | --- |
| CD45.1 (clone A20) | PE | Biolegend | 110707 |
| CD45.2 (clone 104) | PB | Biolegend | 109820 |
| CD45 (clone 30-F11) | PE | Biolegend | 103106 |
| Ly6G (clone 1A8) | FITC | Biolegend | 127606 |
| CD11b (clone M1/70) | BV510 | Biolegend | 101263 |
| MHCII (clone M5/114.15.2) | PE | Biolegend | 107608 |
| NR4A1 (clone 1E10A15) | AF594 | Biolegend | 653504 |
| CD4 (clone 17A2) | FITC | Biolegend | 100204 |
| CD162 (clone 2PH1) | BV421 | BD Horizon | 562807 |
| MYO-1F | AF647 | SAB | C44670 |
| CD11c (clone N418) | VioBlue | Miltenyi | 130-102-797 |
| Gr-1 (clone RB6-(c5) | BV421 | Biolegend | 108433 |
| Zombie Green (clone30-F11) | A700 | Biolegend | 423112 |
| LY6G (clone 1A8) | APC | Biolegend | 127614 |
| CD3 (17A2) | AF700 | Biolegend | 100216 |

## Literature

Amezquita, R. A., Lun, A. T. L., Becht, E., Carey, V. J., Carpp, L. N., Geistlinger, L., Marini, F., Rue-Albrecht, K., Risso, D., Soneson, C., Waldron, L., Pagès, H., Smith, M. L., Huber, W., Morgan, M., Gottardo, R., & Hicks, S. C. (2020). Orchestrating single-cell analysis with Bioconductor. Nature methods, 17(2), 137–145.

Andreatta, M., & Carmona, S. J. (2021). UCell: Robust and scalable single-cell gene signature scoring. Computational and structural biotechnology journal, 19, 3796–3798.

Beuker, C., Schafflick, D., Strecker, JK. et al. Stroke induces disease-specific myeloid cells in the brain parenchyma and pia. Nat Commun 13, 945 (2022).

Beuker, C., Strecker, J. K., Rawal, R., Schmidt-Pogoda, A., Ruck, T., Wiendl, H., Klotz, L., Schäbitz, W. R., Sommer, C. J., Minnerup, H., Meuth, S. G., & Minnerup, J. (2021). Immune Cell Infiltration into the Brain After Ischemic Stroke in Humans Compared to Mice and Rats: a Systematic Review and Meta-Analysis. Translational stroke research, 12(6), 976–990.

Cai, W., Liu, S., Hu, M., Huang, F., Zhu, Q., Qiu, W., Hu, X., Colello, J., Zheng, S. G., & Lu, Z. (2020). Functional Dynamics of Neutrophils After Ischemic Stroke. Translational stroke research, 11(1), 108–121.

Corces, M. R. et al. (2017) ‘An improved ATAC-seq protocol reduces background and enables interrogation of frozen tissues’, Nature Methods. Nature Publishing Group, 14(10), pp. 959–962. doi: 10.1038/nmeth.4396.

Crean, D., & Murphy, E. P. (2021). Targeting NR4A Nuclear Receptors to Control Stromal Cell Inflammation, Metabolism, Angiogenesis, and Tumorigenesis. Frontiers in cell and developmental biology, 9, 589770.

Denorme, F., Portier, I., Rustad, J. L., Cody, M. J., de Araujo, C. V., Hoki, C., Alexander, M. D., Grandhi, R., Dyer, M. R., Neal, M. D., Majersik, J. J., Yost, C. C., & Campbell, R. A. (2022). Neutrophil extracellular traps regulate ischemic stroke brain injury. The Journal of clinical investigation, 132(10), e154225.

Dhanesha, N., Patel, R. B., Doddapattar, P., Ghatge, M., Flora, G. D., Jain, M., Thedens, D., Olalde, H., Kumskova, M., Leira, E. C., & Chauhan, A. K. (2022). PKM2 promotes neutrophil activation and cerebral thromboinflammation: therapeutic implications for ischemic stroke. Blood, 139(8), 1234–1245.

Ermine, C. M., Bivard, A., Parsons, M. W., & Baron, J. C. (2021). The ischemic penumbra: From concept to reality. International journal of stroke : official journal of the International Stroke Society, 16(5), 497–509.

Gao, X., Zhao, X., Li, J., Liu, C., Li, W., Zhao, J., Li, Z., Wang, N., Wang, F., Dong, J., Yan, X., Zhang, J., Hu, X., Jin, J., Mang, G., Ma, R., & Hu, S. (2024). Neutrophil extracellular traps mediated by platelet microvesicles promote thrombosis and brain injury in acute ischemic stroke. Cell communication and signaling : CCS, 22(1), 50.

Gullotta, G. S., De Feo, D., Friebel, E., Semerano, A., Scotti, G. M., Bergamaschi, A., Butti, E., Brambilla, E., Genchi, A., Capotondo, A., Gallizioli, M., Coviello, S., Piccoli, M., Vigo, T., Della Valle, P., Ronchi, P., Comi, G., D’Angelo, A., Maugeri, N., Roveri, L., … Bacigaluppi, M. (2023). Age-induced alterations of granulopoiesis generate atypical neutrophils that aggravate stroke pathology. Nature immunology, 24(6), 925–940.

Jayaraj, R. L., Azimullah, S., Beiram, R., Jalal, F. Y., & Rosenberg, G. A. (2019). Neuroinflammation: friend and foe for ischemic stroke. Journal of neuroinflammation, 16(1), 142.

Jickling, G. C., Liu, D., Ander, B. P., Stamova, B., Zhan, X., & Sharp, F. R. (2015). Targeting neutrophils in ischemic stroke: translational insights from experimental studies. Journal of cerebral blood flow and metabolism : official journal of the International Society of Cerebral Blood Flow and Metabolism, 35(6), 888–901.

Jin, P., Duan, X., Huang, Z., Dong, Y., Zhu, J., Guo, H., Tian, H., Zou, C. G., & Xie, K. (2025). Nuclear receptors in health and disease: signaling pathways, biological functions and pharmaceutical interventions. Signal transduction and targeted therapy, 10(1), 228.

Kao, M. H., Huang, C. Y., Cheung, W. M., Yan, Y. T., Chen, J. J., Ho, Y. S., Hsu, C. Y., & Lin, T. N. (2023). Activating Transcription Factor 3 Diminishes Ischemic Cerebral Infarct and Behavioral Deficit by Downregulating Carboxyl-Terminal Modulator Protein. International journal of molecular sciences, 24(3), 2306.

Kim, M. H., Yang, D., Kim, M., Kim, S. Y., Kim, D., & Kang, S. J. (2017). A late-lineage murine neutrophil precursor population exhibits dynamic changes during demand-adapted granulopoiesis. Scientific reports, 7, 39804. 10.1038/srep39804

Lapostolle, A., Loyer, C., Elhorany, M., Chaigneau, T., Bielle, F., Alamowitch, S., Clarençon, F., & Elbim, C. (2023). Neutrophil extracellular traps in ischemic stroke thrombi are associated with poor clinical outcome. Stroke: Vascular and Interventional Neurology, 3(3).

Lee, S. L., Wesselschmidt, R. L., Linette, G. P., Kanagawa, O., Russell, J. H., & Milbrandt, J. (1995). Unimpaired thymic and peripheral T cell death in mice lacking the nuclear receptor NGFI-B (Nur77). Science (New York, N.Y.), 269(5223), 532–535.

Lefort CT, et al. Distinct roles for talin-1 and kindlin-3 in LFA-1 extension and affinity regulation. Blood. 2012;119(18):4275–4282.

Li, Q., Ye, J., Li, Z., Xiao, Q., Tan, S., Hu, B., & Jin, H. (2024). The role of neutrophils in tPA thrombolysis after stroke: a malicious troublemaker. Frontiers in immunology, 15, 1477669.

Liang, Z., Lou, Y., Hao, Y., Li, H., Feng, J., & Liu, S. (2023). The Relationship of Astrocytes and Microglia with Different Stages of Ischemic Stroke. Current neuropharmacology, 21(12), 2465–2480.

Liebmann, M., Hucke, S., Koch, K., Eschborn, M., Ghelman, J., Chasan, A. I., Glander, S., Schädlich, M., Kuhlencord, M., Daber, N. M., Eveslage, M., Beyer, M., Dietrich, M., Albrecht, P., Stoll, M., Busch, K. B., Wiendl, H., Roth, J., Kuhlmann, T., & Klotz, L. (2018). Nur77 serves as a molecular brake of the metabolic switch during T cell activation to restrict autoimmunity. Proceedings of the National Academy of Sciences of the United States of America, 115(34), E8017–E8026.

Liu, P., Chen, Y., Zhang, Z., Yuan, Z., Sun, J. G., Xia, S., Cao, X., Chen, J., Zhang, C. J., Chen, Y., Zhan, H., Jin, Y., Bao, X., Gu, Y., Zhang, M., & Xu, Y. (2023). Noncanonical contribution of microglial transcription factor NR4A1 to post-stroke recovery through TNF mRNA destabilization. PLoS biology, 21(7), e3002199.

Mazaira, G. I., Zgajnar, N. R., Lotufo, C. M., Daneri-Becerra, C., Sivils, J. C., Soto, O. B., Cox, M. B., & Galigniana, M. D. (2018). The Nuclear Receptor Field: A Historical Overview and Future Challenges. Nuclear receptor research, 5, 101320.

Murphy, E. P., & Crean, D. (2022). NR4A1-3 nuclear receptor activity and immune cell dysregulation in rheumatic diseases. Frontiers in medicine, 9, 874182.

Neumann, J., Sauerzweig, S., Rönicke, R., Gunzer, F., Dinkel, K., Ullrich, O., Gunzer, M., & Reymann, K. G. (2008). Microglia cells protect neurons by direct engulfment of invading neutrophil granulocytes: a new mechanism of CNS immune privilege. The Journal of neuroscience: the official journal of the Society for Neuroscience, 28(23), 5965–5975.

Noll, J. M., Augello, C. J., Kürüm, E., Pan, L., Pavenko, A., Nam, A., & Ford, B. D. (2022). Spatial Analysis of Neural Cell Proteomic Profiles Following Ischemic Stroke in Mice Using High-Plex Digital Spatial Profiling. Molecular neurobiology, 59(12), 7236–7252.

Otxoa-de-Amezaga, A., Miró-Mur, F., Pedragosa, J., Gallizioli, M., Justicia, C., Gaja-Capdevila, N., Ruíz-Jaen, F., Salas-Perdomo, A., Bosch, A., Calvo, M., Márquez-Kisinousky, L., Denes, A., Gunzer, M., & Planas, A. M. (2019). Microglial cell loss after ischemic stroke favors brain neutrophil accumulation. Acta neuropathologica, 137(2), 321–341. h

Pick, R., Begandt, D., Stocker, T. J., Salvermoser, M., Thome, S., Böttcher, R. T., Montanez, E., Harrison, U., Forné, I., Khandoga, A. G., Coletti, R., Weckbach, L. T., Brechtefeld, D., Haas, R., Imhof, A., Massberg, S., Sperandio, M., & Walzog, B. (2017). Coronin 1A, a novel player in integrin biology, controls neutrophil trafficking in innate immunity. Blood, 130(7), 847–858.

Powers, W. J., Rabinstein, A. A., Ackerson, T., Adeoye, O. M., Bambakidis, N. C., Becker, K., Biller, J., Brown, M., Demaerschalk, B. M., Hoh, B., Jauch, E. C., Kidwell, C. S., Leslie-Mazwi, T. M., Ovbiagele, B., Scott, P. A., Sheth, K. N., Southerland, A. M., Summers, D. V., & Tirschwell, D. L. (2019). Guidelines for the Early Management of Patients With Acute Ischemic Stroke: 2019 Update to the 2018 Guidelines for the Early Management of Acute Ischemic Stroke: A Guideline for Healthcare Professionals From the American Heart Association/American Stroke Association. Stroke, 50(12), e344–e418.

Pu, Z. Q., Liu, D., Lobo Mouguegue, H. P. P., Jin, C. W., Sadiq, E., Qin, D. D., Yu, T. F., Zong, C., Chen, J. C., Zhao, R. X., Lin, J. Y., Cheng, J., Yu, X., Li, X., Zhang, Y. C., Liu, Y. T., Guan, Q. B., & Wang, X. D. (2020). NR4A1 counteracts JNK activation incurred by ER stress or ROS in pancreatic β-cells for protection. Journal of cellular and molecular medicine, 24(24), 14171–14183.

R Core Team (2024). R: A Language and Environment for Statistical Computing. R Foundation for Statistical Computing, Vienna, Austria.

Ranhotra H. S. (2015). The NR4A orphan nuclear receptors: mediators in metabolism and diseases. Journal of receptor and signal transduction research, 35(2), 184–188.

Salaudeen, M. A., Bello, N., Danraka, R. N., & Ammani, M. L. (2024). Understanding the Pathophysiology of Ischemic Stroke: The Basis of Current Therapies and Opportunity for New Ones. Biomolecules, 14(3), 305.

Salvermoser, M., Pick, R., Weckbach, L. T., Zehrer, A., Löhr, P., Drechsler, M., Sperandio, M., Soehnlein, O., & Walzog, B. (2018). Myosin 1f is specifically required for neutrophil migration in 3D environments during acute inflammation. Blood, 131(17), 1887–1898.

Sas, A. R., Carbajal, K. S., Jerome, A. D., Menon, R., Yoon, C., Kalinski, A. L., Giger, R. J., & Segal, B. M. (2020). A new neutrophil subset promotes CNS neuron survival and axon regeneration. Nature immunology, 21(12), 1496–1505.

Satija, R., Farrell, J. A., Gennert, D., Schier, A. F., & Regev, A. (2015). Spatial reconstruction of single-cell gene expression data. Nature biotechnology, 33(5), 495–502.

Schindelin, J., Arganda-Carreras, I., Frise, E., Kaynig, V., Longair, M., Pietzsch, T., Preibisch, S., Rueden, C., Saalfeld, S., Schmid, B., Tinevez, J. Y., White, D. J., Hartenstein, V., Eliceiri, K., Tomancak, P., & Cardona, A. (2012). Fiji: an open-source platform for biological-image analysis. Nature methods, 9(7), 676–682.

Schulte, D., Küppers, V., Dartsch, N., Broermann, A., Li, H., Zarbock, A., Kamenyeva, O., Kiefer, F., Khandoga, A., Massberg, S., & Vestweber, D. (2011). Stabilizing the VE-cadherin-catenin complex blocks leukocyte extravasation and vascular permeability. The EMBO journal, 30(20), 4157–4170.

Sheng, M., Weng, Y., Cao, Y., Zhang, C., Lin, Y., & Yu, W. (2023). Caspase 6/NR4A1/SOX9 signaling axis regulates hepatic inflammation and pyroptosis in ischemia-stressed fatty liver. Cell death discovery, 9(1), 106.

Srakočić, S., Gorup, D., Kutlić, D., Petrović, A., Tarabykin, V., & Gajović, S. (2023). Reactivation of corticogenesis-related transcriptional factors BCL11B and SATB2 after ischemic lesion of the adult mouse brain. Scientific reports, 13(1), 8539.

Tsao, C. W., Aday, A. W., Almarzooq, Z. I., Anderson, C. A. M., Arora, P., Avery, C. L., Baker-Smith, C. M., Beaton, A. Z., Boehme, A. K., Buxton, A. E., Commodore-Mensah, Y., Elkind, M. S. V., Evenson, K. R., Eze-Nliam, C., Fugar, S., Generoso, G., Heard, D. G., Hiremath, S., Ho, J. E., Kalani, R., … American Heart Association Council on Epidemiology and Prevention Statistics Committee and Stroke Statistics Subcommittee (2023). Heart Disease and Stroke Statistics-2023 Update: A Report From the American Heart Association. Circulation, 147(8), e93–e621.

Waseem, A., Rashid, S., Rashid, K., Khan, M. A., Khan, R., Haque, R., Seth, P., & Raza, S. S. (2023). Insight into the transcription factors regulating Ischemic stroke and glioma in response to shared stimuli. Seminars in cancer biology, 92, 102–127.

Wickham H., ggplot2: Elegant Graphics for Data Analysis. Springer-Verlag New York, 2016.

Wu, L., & Chen, L. (2018). Characteristics of Nur77 and its ligands as potential anticancer compounds (Review). Molecular medicine reports, 18(6), 4793–4801.

Xiong, Y., Ran, J., Xu, L., Tong, Z., Adel Abdo, M. S., Ma, C., Xu, K., He, Y., Wu, Z., Chen, Z., Hu, P., Jiang, L., Bao, J., Chen, W., & Wu, L. (2020). Reactivation of NR4A1 Restrains Chondrocyte Inflammation and Ameliorates Osteoarthritis in Rats. Frontiers in cell and developmental biology, 8, 158

Yang, L., Chen, H., Guan, L., & Xu, Y. (2022). Sevoflurane Offers Neuroprotection in a Cerebral Ischemia/Reperfusion Injury Rat Model Through the E2F1/EZH2/TIMP2 Regulatory Axis. Molecular neurobiology, 59(4), 2219–2231.

Zehrer, A., Pick, R., Salvermoser, M., Boda, A., Miller, M., Stark, K., Weckbach, L. T., Walzog, B., & Begandt, D. (2018). A Fundamental Role of Myh9 for Neutrophil Migration in Innate Immunity. Journal of immunology (Baltimore, Md. : 1950), 201(6), 1748–1764.

Zenaro, E., Pietronigro, E., Della Bianca, V., Piacentino, G., Marongiu, L., Budui, S., Turano, E., Rossi, B., Angiari, S., Dusi, S., Montresor, A., Carlucci, T., Nanì, S., Tosadori, G., Calciano, L., Catalucci, D., Berton, G., Bonetti, B., & Constantin, G. (2015). Neutrophils promote Alzheimer’s disease-like pathology and cognitive decline via LFA-1 integrin. Nature medicine, 21(8), 880–886.

Zhan, Y., Du, X., Chen, H., Liu, J., Zhao, B., Huang, D., Li, G., Xu, Q., Zhang, M., Weimer, B. C., Chen, D., Cheng, Z., Zhang, L., Li, Q., Li, S., Zheng, Z., Song, S., Huang, Y., Ye, Z., Su, W., … Wu, Q. (2008). Cytosporone B is an agonist for nuclear orphan receptor Nur77. Nature chemical biology, 4(9), 548–556.

Zhang, X., Li, H., Gu, Y. et al. Repair-associated macrophages increase after early-phase microglia attenuation to promote ischemic stroke recovery. Nat Commun 16, 3089 (2025).

Zheng, G. X., Terry, J. M., Belgrader, P., Ryvkin, P., Bent, Z. W., Wilson, R., Ziraldo, S. B., Wheeler, T. D., McDermott, G. P., Zhu, J., Gregory, M. T., Shuga, J., Montesclaros, L., Underwood, J. G., Masquelier, D. A., Nishimura, S. Y., Schnall-Levin, M., Wyatt, P. W., Hindson, C. M., Bharadwaj, R., … Bielas, J. H. (2017). Massively parallel digital transcriptional profiling of single cells. Nature communications, 8, 14049.

Zhao, L., Hu, H., Gustafsson, J. Å., & Zhou, S. (2020). Nuclear Receptors in Cancer Inflammation and Immunity. Trends in immunology, 41(2), 172–185.

Zucha, D., Abaffy, P., Kirdajova, D., Jirak, D., Kubista, M., Anderova, M., & Valihrach, L. (2024). Spatiotemporal transcriptomic map of glial cell response in a mouse model of acute brain ischemia. Proceedings of the National Academy of Sciences of the United States of America, 121(46), e2404203121.

